# SPTLC1 variants associated with childhood onset amyotrophic lateral sclerosis produce distinct sphingolipid signatures through impaired interaction with ORMDL proteins

**DOI:** 10.1101/2022.04.29.490031

**Authors:** Museer A. Lone, Mari J. Aaltonen, Aliza Zidell, Helio F. Pedro, Jonas Alex Morales Saute, Shalett Mathew, Payam Mohassel, Carsten Bonnemann, Eric A. Shoubridge, Thorsten Hornemann

**Affiliations:** Institute of Clinical Chemistry, University Hospital Zurich, University of Zurich, Zurich, Switzerland; Montreal Neurological Institute, McGill University, Montreal, Canada; Department of Human Genetics, McGill University, Montreal, Canada; Center for Genetic and Genomic Medicine, Hackensack University Medical Center, Hackensack, NJ, USA; Center for Genetic and Genomic Medicine, Hackensack University Medical Center, Hackensack Meridian School of Medicine, Hackensack, NJ, USA; Medical Genetics division and Neurology division, Hospital de Clínicas de Porto Alegre, Porto Alegre, Brazil; Graduate Program in Medicine: Medical Sciences, and Internal Medicine Department; Faculdade de Medicina, Universidade Federal do Rio Grande do Sul, Porto Alegre, Brazil; Neuromuscular and Neurogenetic Disorders of Childhood Section, National Institute of Neurological Disorders and Stroke, NIH, Bethesda, MD, USA

**Keywords:** Juvenile Amyotrophic lateral sclerosis (jALS), sphingolipid, serine-palmitoyltransferase (SPT), serine-palmitoyltransferase subunit 1 (SPTLC1)

## Abstract

Amyotrophic lateral sclerosis (ALS) is a progressive neurodegenerative disease affecting motor neurons. Mutations in the *SPTLC1* subunit of serine-palmitoyltransferase (SPT), which catalyzes the first step in the de novo synthesis of sphingolipids cause childhood-onset ALS. SPTLC1-ALS variants map to a transmembrane domain that interacts with ORMDL proteins, negative regulators of SPT activity. We show that ORMDL binding to the holoenzyme complex is impaired in cells expressing pathogenic SPTLC1-ALS alleles, resulting in increased sphingolipid synthesis and a distinct lipid signature. C-terminal SPTLC1 variants cause the peripheral sensory neuropathy HSAN1 due to the synthesis of 1-deoxysphingolipids (1-deoxySLs) that form when SPT metabolizes L-alanine instead of L-serine. Limiting L-serine availability in SPTLC1-ALS expressing cells increased 1-deoxySL and shifted the SL profile from an ALS to an HSAN1-like signature. This effect was corroborated in an SPTLC1-ALS pedigree in which the index patient uniquely presented with an HSAN1 phenotype, increased 1-deoxySL levels, and an L-serine deficiency. These data demonstrate how pathogenic variants in different domains of SPTLC1 give rise to distinct clinical presentations that are nonetheless modifiable by substrate availability.

## Introduction

Amyotrophic lateral sclerosis (ALS) is a progressive, neurodegenerative disease of the lower and upper motor neurons characterized by severe muscle wasting, eventually leading to paralysis and death (1, 2). About 90% of the ALS cases are sporadic, frequently without a recognized heritable factor, while an increasing number of genetic causes account for about 10% of ALS cases with a familial background, and a number of those without a clear family history (3).

Recently, dominant de novo missense and deletion mutations in *SPTLC1* were associated with childhood-onset ALS (4-6). SPTLC1 and SPTLC2 are essential subunits of the enzyme serine-palmitoyltransferase (SPT) which catalyzes the first and the rate-limiting step in the de novo synthesis of sphingolipids (SLs) (Supplemental Fig1A). SPT typically conjugates palmitoyl-CoA with L-serine in a PLP dependent reaction but it can also metabolize L-alanine and glycine under certain conditions, forming an atypical class of toxic 1-deoxysphingolipids (1-deoxySL) that cannot be metabolized to complex SLs or degraded by canonical SL catabolism (7). Variants in the cytoplasmic domains of SPTLC1 and SPTLC2 result in pathologically increased 1-deoxySL levels, causing the Hereditary Sensory and Autonomic Neuropathy type 1 (HSAN1) (8, 9), an autosomal dominant axonopathy, characterized by a progressive sensory loss with variable autonomic involvement.(10-13). Although the majority of reported SPTLC1-HSAN1variants are associated with sensory symptoms, a significant motor involvement was reported for the two variants, SPTLC1-S331F/Y (14).The SPT enzyme complex resides in the endoplasmic reticulum (ER) membrane and at the ER-mitochondrial contact sites (15). SPT is typically composed of two SPTLC1-SPTLC2 dimers which interact with the accessory subunits ssSPTa/b and the regulatory ORMDL proteins ORMDL1, -2 and -3, which are paralogous and functionally redundant proteins (16, 17). ORMDLs interact with the N-terminal transmembrane domain (TMD) of SPTLC1 and act as lipid sensors negatively regulating SPT activity (18-20). All reported SPTLC1-ALS missense variants reside within this TMD (Fig 1A, Supplemental 1B) and one causes an in-frame splice skip of exon 2 that results in a form of SPTLC1 protein completely lacking this TMD (4). Our previous results showed that SPTLC1-ALS variants caused an unregulated synthesis of sphingolipids that did not respond to increasing concentrations of ORMDL3 in an in vitro assay, suggesting that the pathogenic variants could prevent the association of ORMDL proteins with the holoenzyme complex.

**Figure 1.**
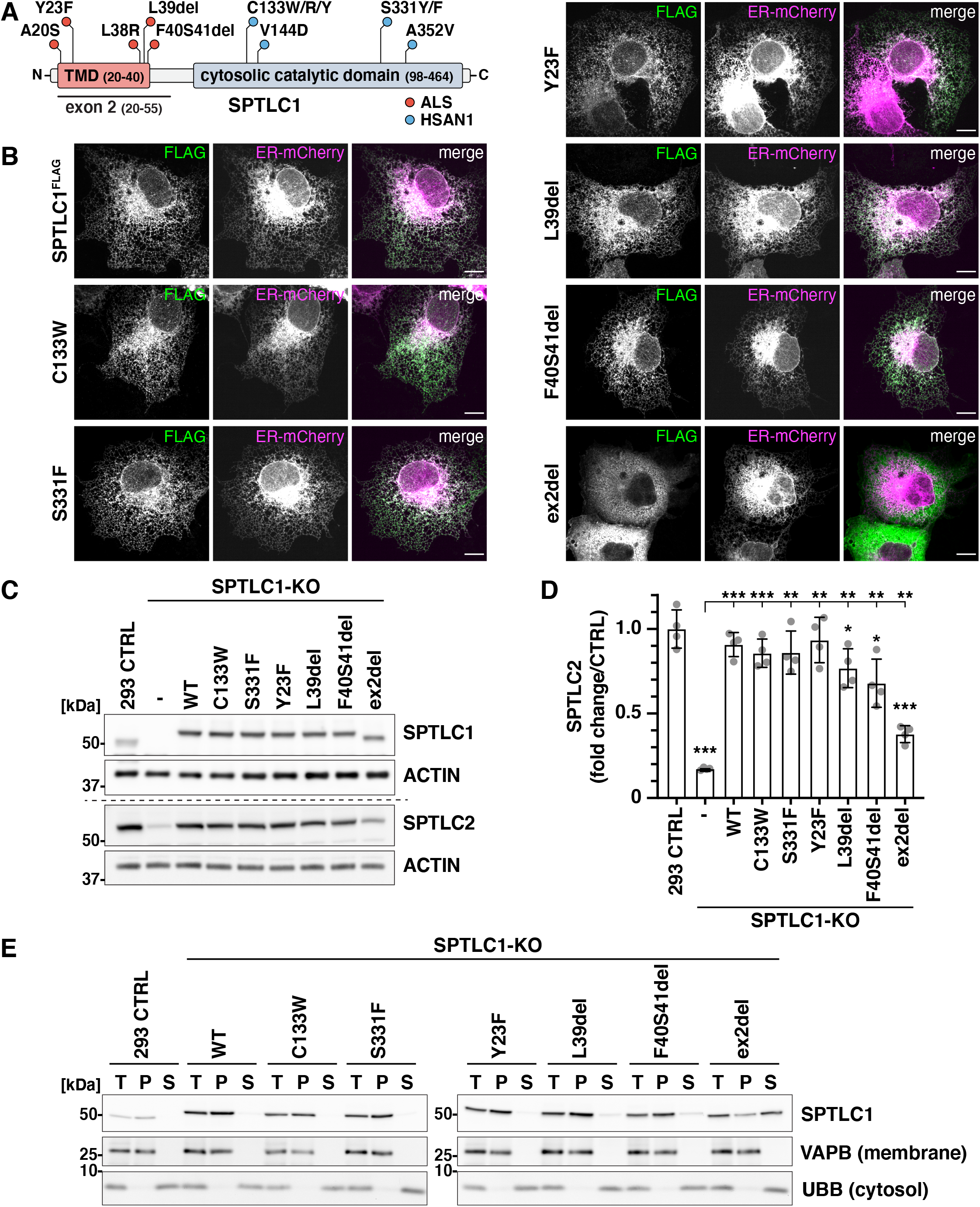
Localization and membrane association of SPTLC1 and pathogenic variants. **(A)** Schematic of SPTLC1 displaying individual protein domains and positions of ALS and HSAN1 pathogenic variants. TMD, transmembrane domain. **(B)** Confocal images of SPTLC1 localization. WT-SPTLC1^FLAG^ and variants were transiently expressed in COS-7 cells and visualized using an anti-FLAG antibody. ER-mCherry serves as an ER marker. Scale bar 10 μm. **(C-D)** Expression of SPTLC1 variants in SPTLC1-KO cells. WT-SPTLC1^FLAG^ and variants were integrated into 293 Flp-In T-REx KO cells and expressed by addition of tetracycline. Whole cell lysates were analyzed by SDS-PAGE and immunoblotting with anti-SPTLC1 and anti-SPTLC2 antibodies (C), and SPTLC2 levels were quantified (D). SPTLC2 signals were normalized to the ACTIN signal. Mean ± SD, n=4 independent replicates, unpaired two-sided Welch’s t-test, * p< 0.05, ** p<0.01, *** p<0.001. **(E)** Analysis of membrane association of SPTLC1 variants. Cell lysates from 293 Flp-In T-Rex control cells and SPTLC1-KO cells expressing WT-SPTLC1^FLAG^ and variants, were centrifuged to separate the membrane pellet and cytosolic supernatant. Equal amounts of total (T), pellet (P) and supernatant (S) were analyzed by SDS-PAGE and immunoblotted with VAPB as a membrane protein control and UBB as a cytosolic protein control.

Here we have expanded the analysis of SPTLC1 variants, tested whether the variants impair the association of ORMDLs with the SPT complex, defined distinct lipid signatures associated with the expression of SPTLC1-ALS and -HSAN1 alleles, and investigated the influence of altered substrate availability on the lipid signatures and clinical phenotypes caused by mutations in different domains of SPTLC1.

## Results

### Loss of exon 2 in *SPTLC1* impairs integration into the ER membrane

SPTLC1 is an ER-localized protein with an N-terminal TMD (Fig 1A, Supplemental Fig 1B). The amino acids in the TMD could be important for the interaction with ORMDL-proteins, but as they could also be essential for ER-targeting and membrane integration of SPTLC1, we first set out to determine the localization and membrane association of a subset of pathogenic SPTLC1 variants in detail, namely the ALS variants Y23F, L39del, F40S41del and a variant missing the whole of exon 2 (ex2del) induced by aberrant splicing in an A20S patient (4) and the HSAN1 variants C133W and S331F (Fig 1A). C-terminally FLAG-tagged SPTLC1 was co-localized with the ER marker Sec61b-mCherry by confocal microscopy when expressed transiently in COS-7 cells, and ER localization was observed for all variants (C133W, S331F, Y23F, L39del and F40S41del), except for the ex2del variant, which lacks the entire TMD and showed a mostly cytosolic distribution when expressed transiently (Fig 1B).

For the biochemical analysis of SPTLC1 variants, C-terminally FLAG-tagged WT-SPTLC1 and C133W, S331F, Y23F, L39del, F40S41del and ex2del variants, were integrated into 293 Flp-In T-REx SPTLC1-KO cells (Fig 1C). The stability of SPTLC2 depends on SPTLC1 (15, 21) and the diminished level of SPTLC2 in SPTLC1-KO cells (17 % of control) was rescued upon re-expression of WT, C133W, S331F and Y23F variants, while a partial rescue was observed by L39del (77 % of control), F40S41del (68 % of control) and ex2del (38 % of control) variants (Fig 1C-D).

Considering that the ex2del variant partially rescues SPTLC2 protein levels, we hypothesized that a fraction of this variant might still be associated with SPTLC2 at the ER and mitochondrial membranes which could have been masked in the microscopy analysis due to a high expression of transiently transfected constructs. To further analyze the membrane association of SPTLC1 and variants, membranes and cytosol were separated by ultracentrifugation from 293 Flp-In T-Rex control cells and variant expressing SPTLC1-KO cells. Endogenous SPTLC1 and WT-SPTLC1^FLAG^ were found in the membrane pellet, as was the ER membrane protein VAPB, while the cytosolic protein UBB was in the supernatant (Fig 1E). All variants were predominantly detected in the membrane pellet fraction, except for the ex2del variant which was enriched in the cytosolic supernatant (Fig 1E). However, part of the ex2del variant was present in the membrane pellet, suggesting that a portion of it is still associated with membranes, likely through interaction of the SPTLC1 aminotransferase domain with SPTLC2.

In summary, neither the HSAN1-or the ALS-causing mutations in SPTLC1 impair the ER localization and membrane association of the protein, except for the TMD lacking ex2del variant which is predominantly soluble in the cytosol and only partially associated with membranes.

### ORMDLs fail to interact with SPTLC1 variants

The patient mutations in *SPTLC1* could impair the interaction of SPTLC1 with ORMDLs, as structural studies have shown that amino acids in the TMD interact with one of the ORMDL-TMDs, and some amino acids close to the catalytic site, such as SPTLC1-S331, interact with the N-terminal loop of ORMDLs that can reach into the active site to occupy the substrate-binding tunnel (16). To analyze the interaction of pathogenic SPTLC1 variants with SPTLC2 and ORMDLs, FLAG-tagged SPTLC1 and variants were purified by FLAG-immunoprecipitation from digitonin-solubilized membrane fractions, and input and eluate fractions were analyzed by immunoblotting. Similar amounts of SPTLC2 co-purified with all variants (Fig 2A); except the ex2del variant which showed a reduced interaction (Fig 2A). However, the level of ex2del variant was lower in the membrane fraction input, as majority of the protein is cytosolic, and these cells also have less SPTLC2 (Fig 1B-D). The interaction with ORMDLs was completely abolished by the ex2del variant which lacks the whole TMD (Fig 2A). The other variants affecting the SPTLC1-TMD, L39del and F40S41del, showed a diminished interaction with ORMDLs, while Y23F retained interaction with ORMDLs (Fig 2A). From the mutations in the catalytic site, both C133W and S331F variants showed a reduced interaction with ORMDLs (Fig 2A).

**Figure 2.**
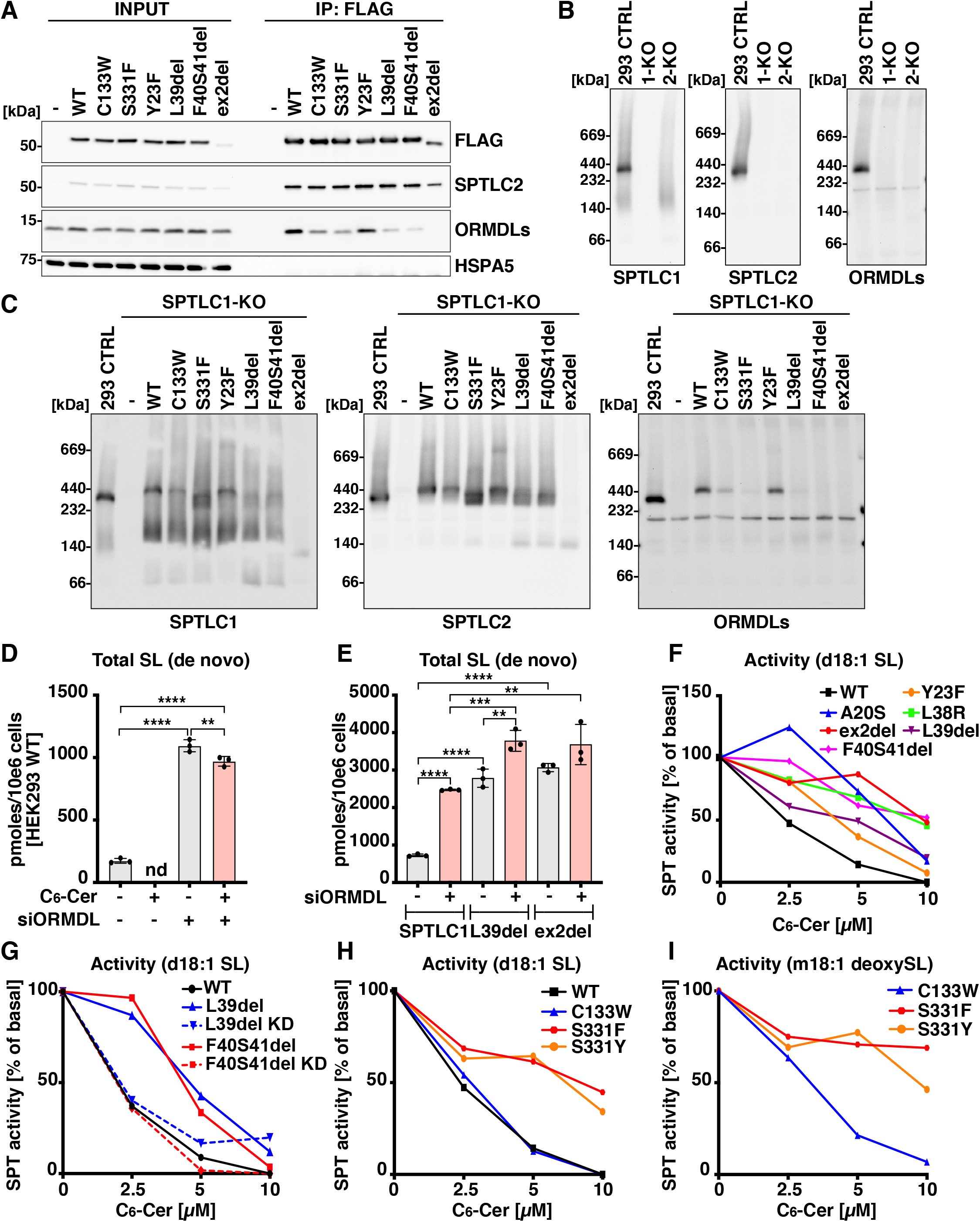
Interaction of SPTLC1 variants with ORMDLs. **(A)** Immunoblot analysis of proteins co-purified with SPTLC1 variants. Membrane fractions from Flp-In T-REx 293 control cells and SPTLC1-KO cells expressing WT-SPTLC1^FLAG^ and variants were solubilized by digitonin, subjected to FLAG-immunoprecipitation. Input (5 %) and eluate (IP:FLAG, 40 %) fractions were analyzed by SDS-PAGE and immunoblotting. HSPA5 was used as a negative control. **(B** and **C)** Analysis of SPT complex by blue native PAGE. Membrane fractions from Flp-In T-REx 293 CTRL, SPTLC1-KO and SPTLC2-KO cells **(B)**, or SPTLC1-KO cells expressing WT-SPTLC1^FLAG^ and variants **(C)** were analyzed by blue native PAGE and immunoblotted with anti-SPTLC1, anti-SPTLC2 and anti-ORMDL antibodies. **(D)** SPT activity in HEK293 WT cells after siRNA mediated knockdown (KD) of ORMDL1-3, in the presence or absence of C_6_-ceramide (C_6_-Cer). **(E)** SPT activity in HEK293 SPTLC1-KO cells expressing WT and indicated SPTLC1-ALS variants after ORMDL1-3 KD. (**F** and **G**) SPT activity after treatment with C_6_-Cer in SPTLC1-KO cells expressing ALS variants **(F)** and in patient derived primary fibroblasts carrying SPTLC1p.L39del and F40S41del mutations (**G**). (**H** and **I**) SPT activity for canonical and 1-deoxySL in SPTLC1-KO cells expressing indicated SPTLC1-HSAN1 variants. SPT activity is measured with incorporation of D_3_, ^15^N-L-serine in SL and D_4_-L-alanine in 1-deoxySL. Mean ± SD, n=3 independent replicates, one-way ANOVA with Bonferroni adjustment for multiple comparisons, ** p<0.01, *** p<0.001, **** p<0.0001.

Next, we asked whether the interaction of ORMDLs with the SPT complex could be detected on a native gel where protein-protein interactions within a protein complex are preserved. The separation of digitonin-solubilized membrane fractions on a native gel, followed by immunoblotting with SPTLC1, SPTLC2 and ORMDL antibodies, showed that ORMDLs migrated together with SPTLC1 and SPTLC2 in a complex of roughly 350∼400-kDa (Fig 2B). This complex was completely absent in SPTLC1-or SPTLC2-KO cells (Fig 2B), while ORMDL-proteins could still be detected on a denaturing gel, although at lower levels compared to control (Supplemental Fig 2A). Notably, the levels of ORMLDs were reduced in SPTLC1-KO (40 % of control) and SPTLC2-KO (49 % of control) cells (Supplemental Fig 2B-C).

To investigate the formation of the SPT complex upon expression of the pathogenic variants, isolated membrane fractions were analyzed on a native gel. Expression of WT-SPTLC1^FLAG^ in SPTLC1-KO cells rescued the formation of SPT-complex as SPTLC1, SPTLC2 and ORMDLs could be detected in a complex, although at a slightly higher molecular weight than in control due to the epitope tag (Fig 2C). In SPTLC1-KO cells expressing S331F, L39del, F40S41del and ex2del variants, the full-size SPT-complex was disassembled as ORMDLs were absent from the complex (Fig 2C), while ORMDLs could still be detected in these samples on a denaturing gel (Supplemental Fig 2D). Instead, S331F, L39del and F40S41del SPTLC1 variants formed a complex with SPTLC2 at a slightly lower molecular weight complexes devoid of ORMDLs, and the ex2del variant only allowed the assembly of a ∼140 kDa SPTLC1-SPTLC2 complex, suggesting that loss of exon 2 also interferes with oligomerization of SPTLC1-SPTLC2 heterodimer (Fig 2C). SPTLC1, SPTLC2 and ORMDLs could still be detected in the full SPT complex with the Y23F and C133W variants, although C133W showed lower levels of this complex (Fig 2C). We observed a disassembly of the SPT complex also in S331Y and L39del patient fibroblasts, with reduced ORMDLs in the SPT complex and a shift in SPTLC2, although the phenotype was not as dramatic as upon expression of variants in a KO background, likely due to patient cells having one wild type allele (Supplemental Fig 2E-F).

In conclusion, co-immunoprecipitation and native PAGE analyses show that pathogenic SPTLC1 variants S331F, L39del, F40S41del and ex2del lose interaction with ORMDLs, leading to a shift in the size of the SPTLC1-SPTLC2 complex. The C133W variant also showed a reduced interaction with ORMDLs, while Y23F supported the interaction and holoenzyme complex assembly.

### SPTLC1 variants show impaired regulation by ORMDLs

We next investigated the enzymatic properties of pathogenic SPTLC1 variants by analyzing de novo SL synthesis in HEK293 SPTLC1-KO cells expressing the pathogenic SPTLC1 variants. Cellular SPT activity was analyzed with a metabolic labelling assay, in which cells were grown in serine and alanine deficient medium supplemented with stable isotope labelled D_3_-^15^N-L-serine (to label canonical SL) and D_4_-L-alanine (to label 1-deoxySL). For quantification, total sphingolipids were extracted and quantified by high-resolution mass spectrometry. The interaction of ORMDLs with SPT inhibits SPT activity, as knockdown (Fig 2D), (22) or knockout (23) of all three ORMDLs leads to increased SL synthesis. In contrast, while the expression of SPTLC1 variants L39del and ex2del in SPTLC1-KO cells led to an overall increase in SL synthesis compared to WT-SPTLC1 expressing cells, silencing ORMDLs had little or no effect on the activity of the L39del and ex2del variant (Fig 2E), suggesting that increased SL synthesis by the variants is primarily caused by dysfunctional feedback inhibition rather than by an increased activity of the holoenzyme.

ORMDLs play a role in homeostatic feedback regulation of SPT activity by ceramides (24), as addition of cell permeable C_6_-ceramide (C_6_-Cer) inhibits SL synthesis while the inhibitory effect is blunted in the absence of ORMDLs (Fig 2D) (22). In WT-SPTLC1 expressing cells, de novo SL synthesis was gradually reduced and ultimately suppressed with increasing concentrations of C_6_-Cer (Fig 2F). In contrast, SPTLC1 variants affecting the TMD showed a reduced response to inhibition of SL synthesis by C_6_-Cer, with the strongest effect seen for the L38R, F40S41del and ex2del variants. An attenuated C_6_-Cer response was also observed in fibroblasts derived from L39del and F40S41del patients, which could be restored after knockdown of the mutant mRNA transcripts (Fig 2G), using siRNAs that specifically silence the expression of the respective mutant alleles as shown previously (4). Additionally, the SL synthesis by variants S331F and S331Y was resistant to C_6_-Cer treatment whereas the response in the C133W expressing cells was similar to WT (Fig 2H). C_6_-Cer induced inhibition was also seen for synthesis of 1-deoxySL in C133W but not in S3331Y and S331F expressing cells (Fig 2I).

In summary, the lack of interaction of SPTLC1-ALS variants with the ORMDLs results in dysfunctional feedback regulation and increased SPT activity, leading to higher SL levels in cells.

### Lipid signatures of motor and sensory neuropathy

To define and compare the distinct alterations in SL classes and species induced in SPTLC1 associated ALS and HSAN1 disease conditions, we analyzed SL profiles in variant expressing cells and patient derived plasma and fibroblasts.

First, we expressed the variants in HEK293 SPTLC1-KO cells and quantified the labeled de novo synthesized SL species, including ceramides, sphingomyelins and 1-deoxyceramides. Compared to WT-SPTLC1 expressing cells, SPTLC1-ALS variants affecting the TMD (A20S, Y23F, L38R, L39del, F40S41del, ex2del) showed an increased formation of canonical SLs, including those with saturated (d18:0), mono-(d18:1) and di-unsaturated (d18:2) sphingoid bases (Fig 3A-F). The highest levels were measured for the ex2del variant, followed by L38R, L39del and F40S41del variants, while A20S and Y23F variants showed modest accumulations. The HSAN1 variant C133W did not increase synthesis of canonical SLs but in contrast induced the formation of 1-deoxySLs (m18:0, m18:1) which were not formed by the TMD variants (Fig 3G-H). The S331F and S331Y variants gave rise to a combined biochemical phenotype with increased synthesis of both canonical and 1-deoxySLs.

**Figure 3.**
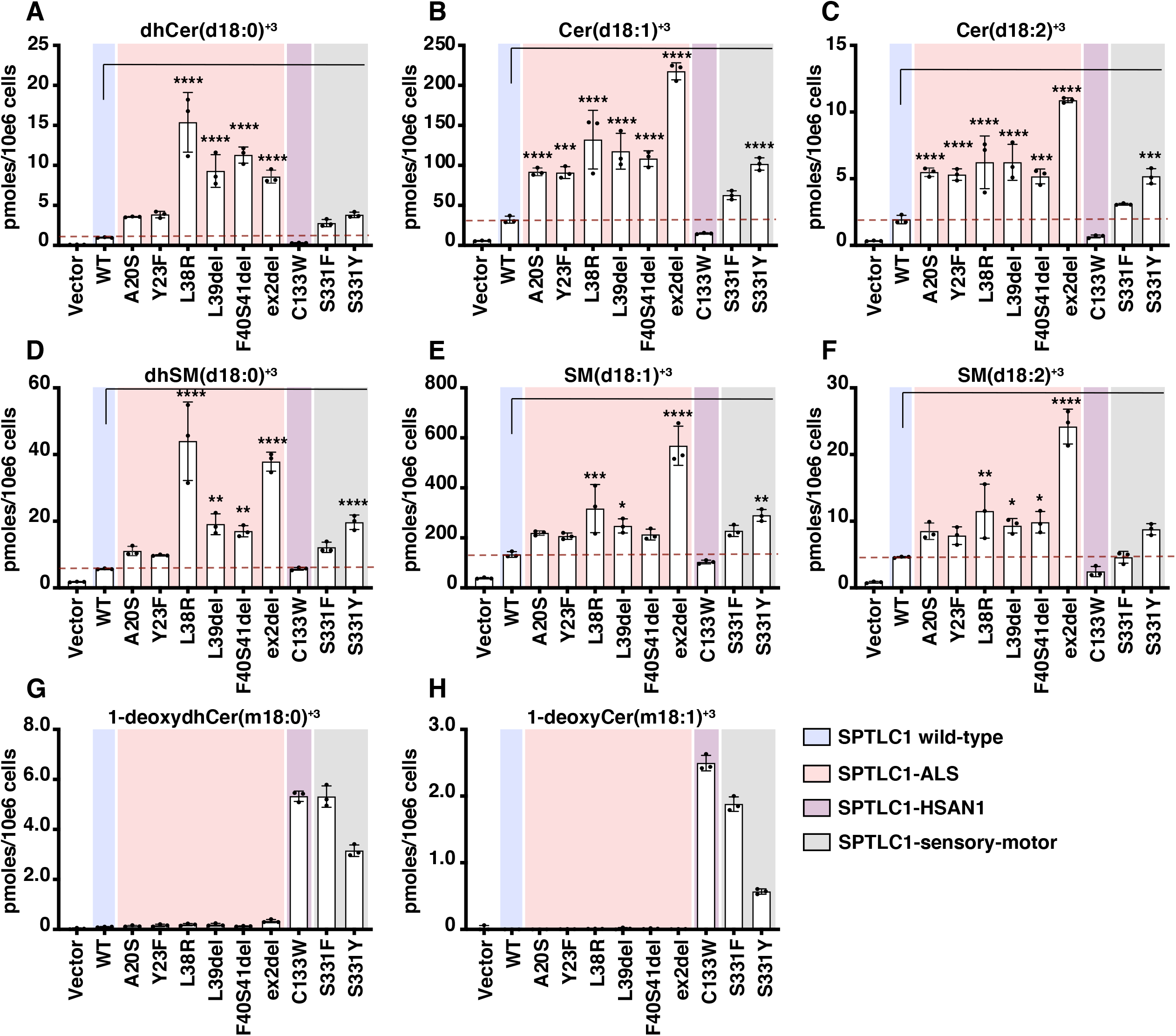
De novo SL synthesis in variant expressing cells. (**A-F**) De novo formation of ceramides (Cer, **A-C**), sphingomyelins (SM, **D-F**), and 1-deoxyceramides (**G** and **H**) in HEK293 SPTLC1-KO cells expressing WT-SPTLC1 and variants. Cells were probed for de novo SPT activity by stable isotope labelling assay, that incorporates D_3_, ^15^N-L-serine in SL and D_4_-L-alanine in 1-deoxySL, inducing a mass shift (+3 Da) in the respective de novo formed SL and absolute levels of each SL species were measured relative to internal lipid standard. Data are represented as mean ± SD, n=3 independent replicates, one-way ANOVA with Bonferroni adjustment for multiple comparison, ** p<0.01, *** p<0.001, **** p<0.0001.

A comparison of the plasma SL profile between the ALS patients carrying Y23F, L39del and F40S41del variants and unrelated HSAN1 patients carrying the C133W variant revealed similar changes in SL profiles, with increased SL levels in ALS patient plasma and higher 1-deoxyceramide levels in the HSAN1 patients (Supplemental Fig 3A-D). There were, however, variations in the SL lipid classes and species, as ceramides, sphingomyelins and hexosyl-ceramides were differently affected in different patients, and the relative increases were more prominent for ceramides with a saturated (d18:0) sphingoid bases compared to unsaturated (d18:1 or d18:2) backbones (Supplemental Fig 3A-D).

To characterize the SL pattern associated with ALS- and HSAN1-SPTLC1 variants, we compared the profile of de novo formed SL species between HEK293 SPTLC1-KO cells expressing the pathogenic SPTLC1 variants. A heatmap cluster analysis showed significant differences in the de novo formed SL profiles (Fig 4A). The HSAN1 variants C133W, S331F and S331Y clustered due to the increase in 1-deoxySL species, while the ALS TMD variants clustered due to the relative enrichment in canonical SL species. The S331 variants showed a mixed pattern which partly resembled the TMD variants due to increased synthesis of canonical SLs, with the S331F variant showing more similarity with the HSAN1 C133W variant, and S331Y with the ALS TMD variants (Fig 4A). The ex2del variant showed the strongest relative accumulation of canonical SL species, followed by L38R, F40S41del, and L39del variants.

**Figure 4.**
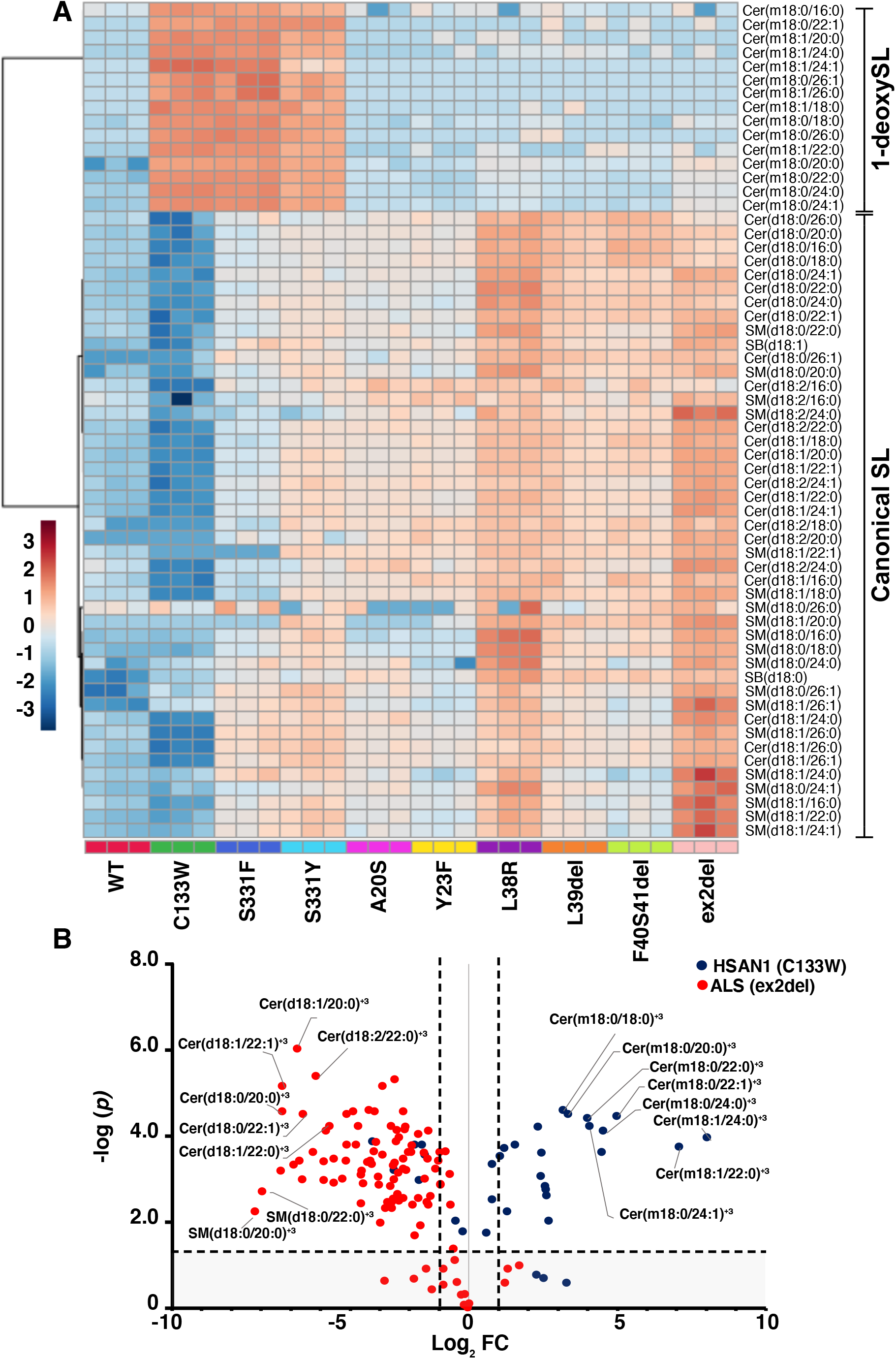
Sphingolipid signatures in variant expressing cells. **(A)** Heatmap cluster analysis of de novo formed SL species from HEK293 SPTLC1-KO expressing WT-, ALS-or HSAN1-SPTLC1 variants. Absolute levels of each SL species were measured relative to internal lipid standard. Shown is plot of the Log transformed (base 10) data with Euclidean distance measure. **(B)** Volcano plot comparing de novo formed SL species in SPTLC1-KO cells expressing the SPTLC1-HSAN1 (C133W) and SPTLC1-ALS (ex2del) variants. MetaboAnalyst Suite 5.0 was used for comparison of species profiles as heat maps and volcano plot. n=3 independent biological replicates, significance (*p*) and fold change (FC) are represented as dotted lines.

We next explored the changes in SL species in detail by volcano plot comparison of SL enrichment in HSAN1 variant C133W expressing cells relative to ALS variant ex2del expressing cells. In addition to the prominent enrichment of de novo formed 1-deoxyceramide species in C133W expressing cells, in ex2del expressing cells there was an enrichment of distinct ceramide species with uncommon N-acyl-chains (Fig 4B). The most abundant ceramides in HEK293 cells carry fatty acids C_16_, C_24:0_ and C_24:1_, while ceramides with C_18:0_, C_20:0_, C_22:0_ and C_22:1_ acyl chains, which have very low abundance in WT-expressing cells, accumulated in cells expressing ex2del (Fig 4B) and also in cells expressing other TMD and S331 variants (Supplemental Fig 4A-B). A similar acyl-chain pattern was seen in primary fibroblasts isolated from L39del and F40S41del patients (Supplemental Fig 5A-B) in which targeted siRNA mediated knockdown of the mutant alleles normalized the acyl-chain profiles (Supplemental Fig 5C). Lastly, the plasma species profiles in patients carrying Y23F, L39del, and F40S41del showed similar changes in ceramide species (Supplemental Fig 6).

In conclusion, the lipid signature for ALS associated variants is characterized by increased SL synthesis and accumulation of uncommon acyl-chain length ceramide species, whereas in HSAN1 the signature is characterized by increased deoxySL synthesis.

### Serine availability modulates the clinical presentation of SPTLC1 variants

SL analysis showed clear differences between the SPTLC1 variants when expressed in cells; however, the correlation of variants with disease presentation is not always straightforward, and L39del and S331 variants have been reported to cause both sensory and motor phenotypes (4, 25) suggesting that additional factors influence disease outcome.

L-serine and L-alanine availability modulate the synthesis of canonical and 1-deoxy lipids, respectively, as 1-deoxySL formation is induced under conditions of L-serine restriction (26, 27). We therefore tested the effect of reduced serine availability on sphingolipid profiles of SPTLC1 variant expressing cells. WT-SPTLC1, ex2del and L39del expressing cells were cultured at decreasing L-serine and constant L-alanine concentrations. In the absence of L-serine, we observed a reduction in total SL synthesis (Fig 5A) and an increase in 1-deoxySL synthesis in both WT and variant expressing cells (Fig 5B). However, the extent of 1-deoxySL formation with reduced serine conditions was higher in variant expressing cells than in those expressing WT-SPTLC1. Interestingly, the L39del variant showed a significantly higher 1-deoxySL accumulation than the ex2del under these conditions. (Fig 5B). Heatmap cluster analysis showed a gradual shift of SL enrichment profiles from the ALS lipid signature toward the HSAN1 signature which is distinguishable for the L39del mutant expressing cells (Fig 5C).

**Figure 5.**
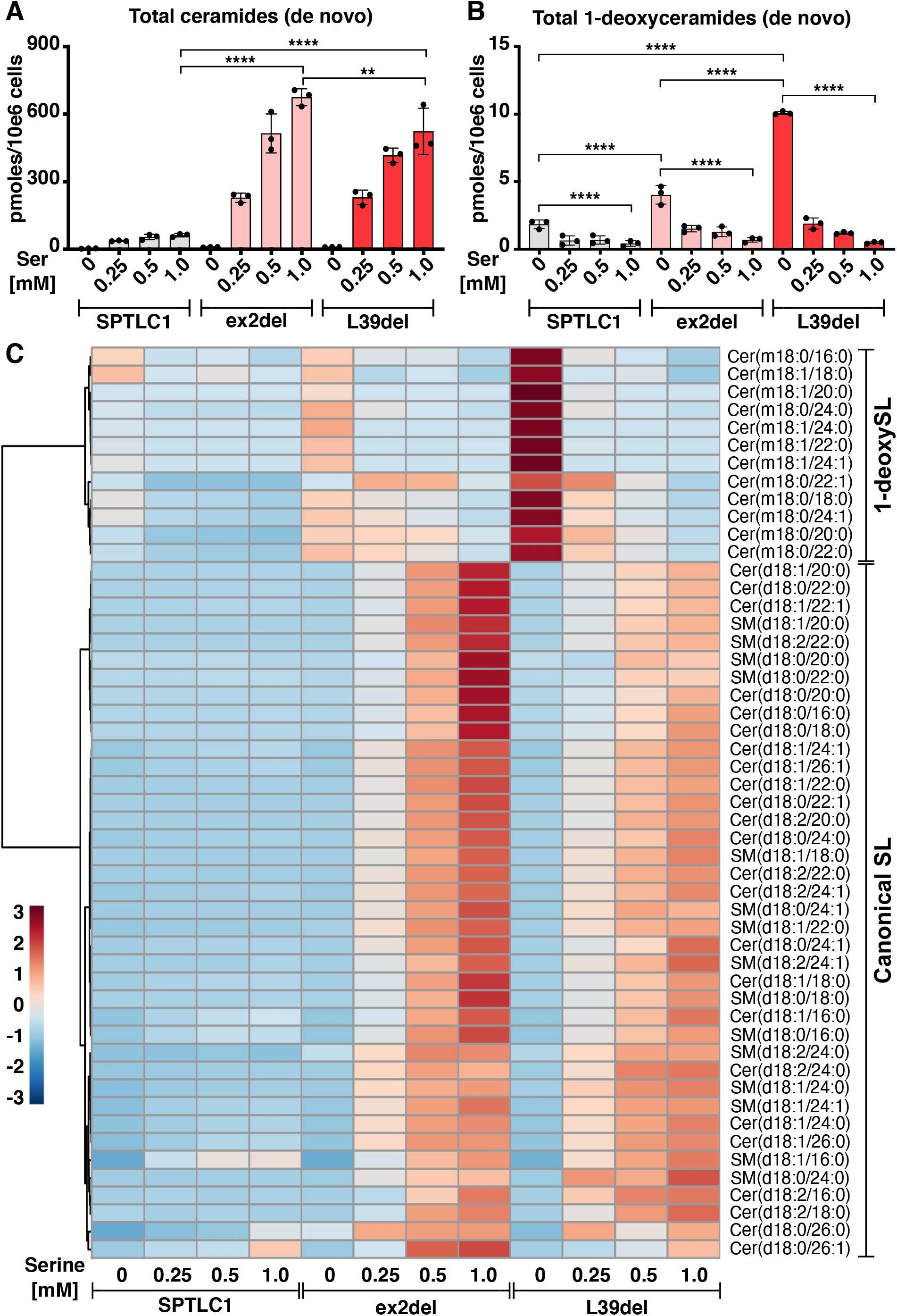
Sphingolipid signatures shift upon amino acid availability. (**A** and **B**) Total de novo formed (**A**) ceramides and (**B**) 1-deoxyceramides in HEK293 SPTLC1-KO cells expressing SPTLC1 WT, ex2del, and L39del variants. Cells were treated with increasing D_3_,^15^N-L-serine (0 to 1 mM) in the presence of a constant D_4_-alanine concentration (2 mM). Data are represented as mean ± SD, n=3 independent replicates, one-way ANOVA with Bonferroni adjustment was used for pair wise comparison ** p<0.01, **** p<0.0001. **(C)** Heatmap cluster analysis of averaged SL species as in A and B. Log transformed (base 10) data with Euclidean distance measure were plotted using MetaboAnalyst Suite 5.0.

The modulation of SL synthesis by substrate availability suggests that the concentration of L-serine and L-alanine in patients might influence the clinical presentation of SPTLC1 variants. We investigated this in the previously reported L39del family (4) where four children (II-1, II-2, II-3 and II-4) showed an ALS disease phenotype, and the father (I-1) presented with a sensory neuropathy that was initially diagnosed as HSAN1 (Fig 6A). The difference in the clinical presentation was reflected at plasma lipid levels which revealed elevated 1-deoxySLs for the father (I-1) but not for the children (II-1, II-2, II-3 and II-4) or the unaffected mother (I-2) (Fig 6B).

**Figure 6.**
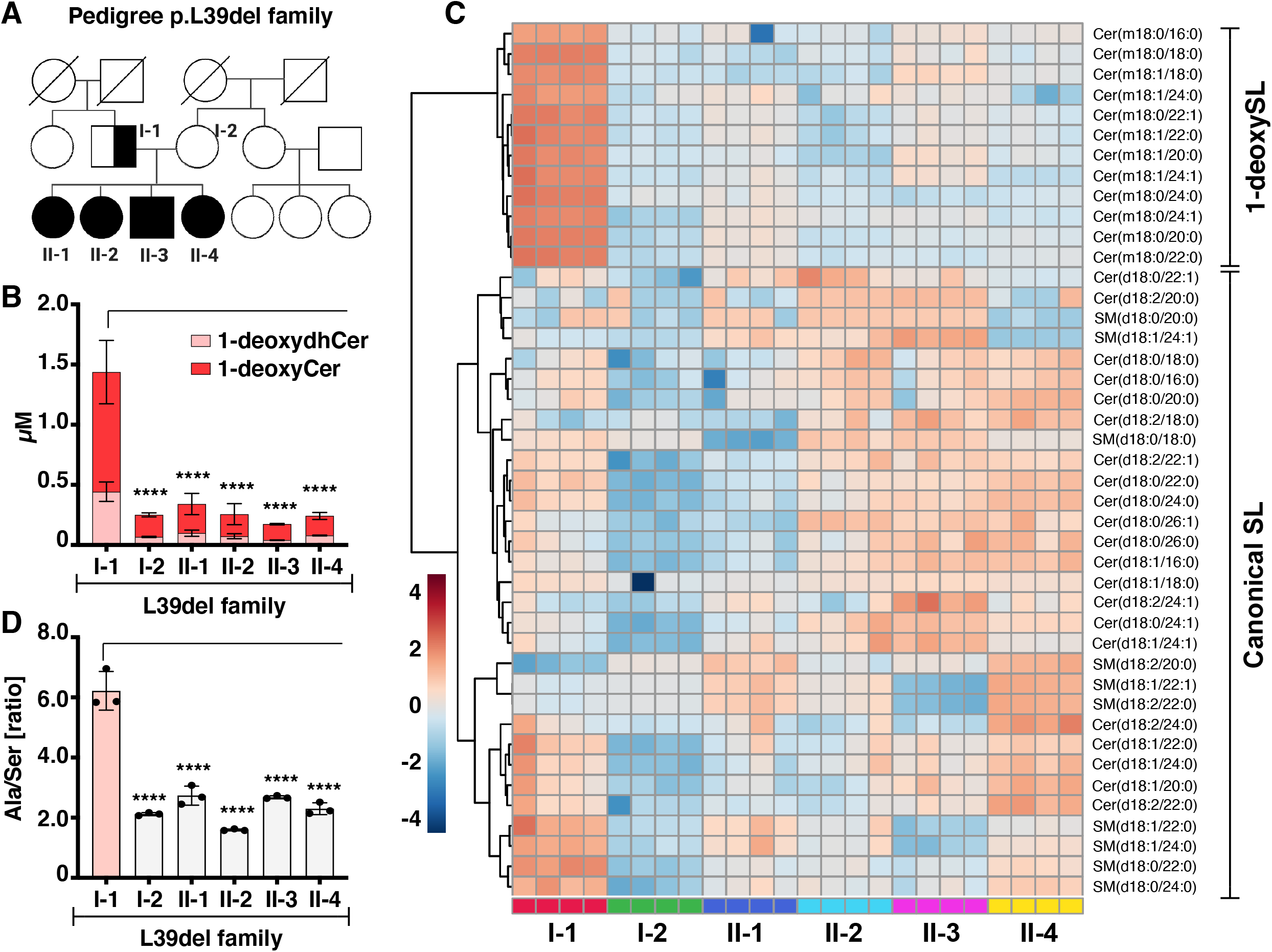
Sphingolipid profiles are influenced by alanine and serine ratios in vivo in the L39del family. **(A)** Pedigree of the family harboring the SPTLC1p.Leu39del mutation. Half-filled shapes represents a sensory phenotype, filled shapes reflect members with motor neuron disease. Circles= females, boxes= males, I and II indicate first and second generation individuals. **(B)** Total plasma 1-deoxydihydroceramides (m18:0) and 1-deoxyceramides (m18:1) in individual family members. Data are represented as mean ± SD, n=4, two-way ANOVA with Dunnett’s adjustment for multiple comparisons. (**C**). Heatmap cluster analysis of SL species in the plasma of the family members. Lipids were extracted and quantified from plasma independently 4 times. Log transformed (base 10) data with Euclidean distance measure were plotted using MetaboAnalyst Suite 5.0. (**D**) Alanine/serine ratio in the plasma of individual family members. Data are represented as mean ± SD, n=3, one-way ANOVA with Bonferroni adjustment for multiple comparison, **** p<0.0001.

A heatmap cluster analysis of plasma SLs within the family (Fig 6C) revealed that the lipid signature of the father resembled the pattern of HSAN1 variant C133W expressing cells (Fig 4A) but the children had a lipid signature pattern similar to cells expressing L39del or other TMD variants (Fig 4A). Since the L-serine and L-alanine concentrations are able to modulate SL and 1-deoxySL enrichment in L39del expressing cells (Fig 5A-B), we wanted to test whether altered amino acid balance could explain the SL differences in patients. Notably, the father showed an increased ratio of L-alanine to L-serine compared to the family members (Fig 6D) which was primarily driven by father’s low plasma L-serine levels (Supplemental Fig 7A) as L-alanine levels were similar (Supplemental Fig 7B).

In summary, in addition to the genetic variations in SPTLC1, the availability of L-serine and L-alanine in patients affects the clinical phenotype and can shift the lipid signature between HSAN1 and ALS.

## Discussion

This study demonstrates that pathogenic variants in different domains of the SPTLC1 subunit of SPT generate distinct lipid signatures that are associated with specific clinical phenotypes. The variants associated with childhood onset ALS cluster in the TMD, while those in the cytosolic domain are largely associated with HSAN1. The transmembrane pathogenic variants do not prevent localization to the ER, except in the case where the TMD is deleted, but they generally impair binding of the negative regulator ORMDLs to the holoenzyme complex, resulting in increased SL synthesis and distinct lipid signatures.

The SPTLC1-ALS variants in the TMD: ex2del, F40S41del and L39del, showed the least interaction with ORMDLs (Fig 2A, C) and, together with the adjacent L38R variant, showed impaired feedback inhibition by C_6_-Cer (Fig 2F), causing a large increase in SL synthesis (Fig 3), similar to cells in which all three ORMDLs were silenced (Fig 2D-E). The ex2del variant showed highest SL synthesis despite its mis-localization and reduced interaction with SPTLC2, highlighting the importance of SPTLC1-TMD in regulating SPT activity. In contrast, Y23F retained the interaction with ORMDLs and together with A20S, only caused a modest increase in SL levels, suggesting that changes in the luminal residues of the TMD (Supplemental Fig 1B) are better tolerated. Additional ORMDL-independent homeostatic control points in the pathway acting downstream of SPT could exist that are not affected by the variants.

We observed differences in the saturation of sphingoid backbones (Fig 3), and the conjugated N-acyl chains (Fig 4, Supplemental Fig 4, Supplemental Fig 5, and Supplemental Fig 6) of SL species formed by the variants. These differences are not easily explained by an increased SPT activity as most of these modifications occur downstream of SPT. N-acylation of the spingoid backbone is determined by CERS enzymes (28) while the double-bonds in the SL backbones are introduced by the desaturases DEGS1 and FADS3 (Supplemental Fig 1A) (29, 30). In the SPTLC1-ALS expressing cells and patient derived fibroblasts, the relative increase was the highest for SLs conjugated to C_18:0_, C_20:0_, C_22:0_ acyl chains, which are usually minor species and barely detected in control cells (Fig 4, Supplemental Fig 4 and Supplemental Fig 5). A possible explanation, at least in the case of the largely cytosolic ex2del variant, might be the distinct intracellular localization of the variant, which might affect local concentration of the metabolic intermediates and the reaction kinetics of the subsequent enzymes in the pathway. In any case, these intermediate chain length species specifically increased in the SPTLC1-ALS could serve as potential biomarkers for this rare condition (Supplemental Fig 6) and could also be the underlying cause for the specific motor neuron toxicity in SPTLC1-ALS.

The SPTLC1-ALS variants do not typically form 1-deoxySLs, which are the hallmark for the SPTLC1-HSAN1 variants. The A20S variant was previously shown to mediate 1-deoxySL formation in vitro (31), however these results were not confirmed in our cell based assays and under these conditions (Fig 3G and H). SPTLC1-HSAN1 variants S331F and S331Y showed a mixed SL phenotype that was characterized by both an increased canonical activity and a significant formation of 1-deoxySLs (Fig 3A to H). This mixed metabolic pattern is mirrored by the clinical presentation of the HSAN1 patients having both sensory and motor symptoms (25, 32-34) Initially, the disease phenotype in the patients was characterized as hereditary motor and sensory neuropathy but the diagnosis was revised to HSAN1 when the sensory symptoms became manifest. Structural studies showed the interaction of S331 with the N-terminal ORMDL-loop (16) and S331 is required for interaction with ORMDLs and for feedback inhibition by ceramides (Fig 2A, C, H), explaining the increased SL accumulation. Surprisingly, the HSAN1 variant C133W also showed reduced association with ORMDLs although the feedback inhibition by C_6_-Cer or synthesis of canonical SLs was similar to WT (Fig 2A, C, and H), indicating that mechanisms other than the binding of ORMDL proteins to the holoenzyme complex may be important in regulating SPT activity.

A heatmap cluster analysis of cells expressing the SPTLC1-ALS and -HSAN1 variants, showed a continuous shift in the lipid signature from the SPTLC1-HSAN1 to the SPTLC1-ALS variants (Fig 4A). On one end of the spectrum is the HSAN1 C133W variant that is associated with sensory symptoms and increased 1-deoxySL whereas at the other end the exon2del variant is associated with a motor neuron disease and increase in canonical SL. This pattern suggests that the increased formation of canonical SLs with uncommon N-acyl chains is primarily toxic for the cerebral motor neurons whereas increased 1-deoxySLs primarily affects peripheral sensory neurons. What renders these neuronal populations more susceptible to one or the other SL metabolite remains unknown.

SPTLC1-ALS variants are capable of forming significant amounts of deoxySL in conditions of L-serine deficiency (Fig 5B), and the comparison of the metabolic signatures showed that L-serine deficiency renders the SL pattern in the SPTLC1-ALS expressing cells similar to that of the SPTLC1-HSAN1 expressing cells (Fig 5C). This raises the possibility that the motor phenotype in ALS might be converted to a more HSAN1 like phenotype through changes in amino acid availability, hypothesis that we investigated in the previously reported SPTLC1 L39del family (Fig 6A). Although all affected family members were carriers of the L39del variant, the index patient (I-1) showed an isolated sensory phenotype, initially characterized as HSAN1, with elevated plasma 1-deoxySLs and higher ratio of alanine to serine due to low plasma serine levels (Fig 6B, C and D and Supplemental Fig 7A), compared to the other family members with motor neuron disease. It is currently unclear why the L-serine levels are low in the index case, but it demonstrates that the two disease phenotypes are modifiable by the L-serine/L-alanine ratio. In that respect, the HSAN1 and ALS phenotypes reflect the opposite ends of a spectrum characterized by a variable degree of sensory and motor symptoms depending on the underlying pattern of increased 1-deoxySL versus canonical SL. This has therapeutic consequences as supplementing L-serine in HSAN1 patients should suppress 1-deoxySL formation and slow disease progression, whereas L-serine supplemention in SPTLC1-ALS patients might worsen the phenotype by stimulating the synthesis of canonical SLs.

## Materials and methods

### Cell culture

Cell lines were grown in high-glucose Dulbecco’s Medium (DMEM, Wisent 319-027-CL for interaction analysis and protein biochemistry; or DMEM, D5796 Sigma-Aldrich, St. Louis, MO, USA for lipidomics) supplemented with 10% fetal bovine serum (FBS) in 5% CO_2_ incubator at 37°C. Cell lines were regularly tested for mycoplasma contamination. For immunofluorescence analysis, cells were cultured on glass coverslips.

### Cell lines

For protein analysis, Flp-In T-REx 293 cells (Invitrogen) were used for expressing of FLAG-tagged SPTLC1 variants for the analysis of protein interactions on SDS-PAGE and native PAGE. The generation of HEK293-SPTLC1 knockout cell line (for lipidomics) was reported earlier (4). SPTLC1-KO and SPTLC2-KO Flp-In T-Rex 293 cell lines (protein analysis) were reported before (15). Patient fibroblasts were cultured in DMEM with 10% FBS as reported previously (4).

### Generation of cell lines

To express FLAG-tagged SPTLC1 and pathogenic variants in SPTLC1-KO cells, cells were rescued by integration of tetracycline inducible SPTLC1^FLAG^ on pDEST-pcDNA5 plasmids as described earlier (35). Protein expression was induced with 1 µg/ml tetracycline for 4 hours. Plasmid transfections were performed with lipofectamine 3000 (Thermo Fisher Scientific) in HEK293 SPTLC1-KO cells. Transgenic HEK293 cell lines were selected for growth in DMEM media containing 400 µg/ml Geneticin (Thermo Fischer Scientific) and 1 % penicillin/streptomycin (P/S).

For blue-native PAGE analysis, control and patient fibroblasts were immortalized by transduction with retroviral vectors expressing the HPV-16 E7 gene and the catalytic component of human telomerase (htert), as described previously (36)

### Plasmids

pcDNA5-pDEST-SPTLC1^FLAG^ plasmid was generated by flipping SPTLC1 from pDONR221 donor vector into pcDNA5-pDEST-3xFLAG destination vector (15) using Gateway cloning technology, yielding SPTLC1 with C-terminal triple FLAG-tag. SPTLC1 variants were generated using mutagenesis PCR (QuikChange Lightning, Agilent) in pDONR221-SPTLC1 and flipped into pcDNA5-pDEST-3xFLAG destination vector.

ER-mCherry (mCh-Sec61 beta) plasmid was a gift from Gia Voeltz (Addgene 49155) (37). For lipid analysis, SPTLC1 was cloned into the pCDNA3.1-V5-His-tag vector. PCR based site directed mutagenesis was used to introduce SPTLC1-ALS and -HSAN1 mutations in the plasmid vector backbone (4, 14, 21).

### Confocal microscopy

For confocal microscopy, COS-7 cells were transfected using Lipofectamine 3000 with the indicated pcDNA5-SPTLC1^FLAG^ and ER-marker plasmids, fixed the day after with 6% formaldehyde in PBS, solubilized with 0.1% Triton X-100 in PBS for 15 min at RT and blocked with 5% (w/v) BSA in PBS. Primary and secondary antibodies were diluted in 5% (wt/vol) BSA in PBS, and coverslips were incubated with the antibodies for 1 hour at RT with five washes with PBS in between. Image acquisition (z-stacks, step size 0.2 µm) was performed with an Olympus IX83 confocal microscope containing a spinning disk system (Andor/Yokogawa CSU-X) using Olympus UPLSAPO 100x oil objectives. Images were processed in Fiji.

### Membrane isolation

For separation of membrane and cytosolic fractions, cells were resuspended in fractionation buffer (20 mM HEPES-KOH pH 7.6, 1 mM EDTA, 1x cOmplete protease inhibitor (Roche 11873580001)), lysed by passing through a 25 G needle ten times and unbroken cells were pelleted 5 min at 800g three times. Protein concentration in lysates was determined by Bradford assay, diluted to 1 µg/µl and input sample was collected and mixed with Laemmli buffer. To pellet membranes, 100 µg of lysates were centrifuged 90 min at 100,000g in TLA100 rotor, and soluble fractions were collected in the supernatant and mixed with Laemmli buffer. Membrane pellet was solubilized in Laemmli buffer and equal amounts of each fraction were analyzed by SDS-PAGE and immunoblot.

For preparation of 20,000g membrane fractions for co-immunoprecipitation and BN-PAGE analysis, cells were resuspended in isolation buffer (220 mM mannitol, 70 mM sucrose, 20 mM HEPES-KOH pH7.6, 1 mM EDTA, 1x cOmplete protease inhibitor) and homogenized by 15-20 strokes in a rotating Teflon-glass homogenizer at 1000 rpm. Homogenates were centrifuged twice at 800g for 5 min to remove nuclei and cell debris. Membranes were pelleted at 20,000g for 30 min, resuspended in isolation buffer and protein concentration was determined by Bradford assay.

### Denaturing and native PAGE

For analysis of protein levels in whole cell lysates, cells were lysed with RIPA lysis buffer (1% Triton X-100, 0.1% (w/v) SDS, 0.5% (w/v) sodium deoxycholate, 1 mM EDTA, 50 mM TRIS pH 7.4, 150 mM NaCl, 1x cOmplete protease inhibitor) for 20 min and centrifuged for 15 min at 20 000g. Equal amounts of protein were analyzed on 10% Tris-Tricine SDS-PAGE system (38) with Precision plus protein standard (Bio-Rad 1610363) as a molecular weight marker, and blotted onto a nitrocellulose membrane. Protein intensities were quantified in Fiji.

For analysis of SPT complex on blue-native PAGE, 20,000g membrane fractions were solubilized with 4 g digitonin/g protein (1% wt/vol) for 20 min, centrifuged for 20 min at 20 000g and equal amounts of supernatants were separated on a 4-13% gradient gel as previously described (39) with Amersham HMW native marker kit (17044501 Cytiva) as a molecular weight marker, and blotted onto a PVDF membrane. For SDS-PAGE analysis of protein levels in native samples, equal amounts of supernatants were mixed with Laemmli buffer and ran on SDS-PAGE gel as above.

### FLAG immunoprecipitation

Membrane fractions were solubilized in buffer (50mM TRIS pH 7.4, 150 mM NaCl, 20 mM MgCl_2_, 4 g digitonin/g protein (1% wt/vol), 1x cOmplete protease inhibitor) for 20 min and aggregates were pelleted at 20 000g for 20 min. Input sample was taken from the supernatant and the rest was incubated with anti-FLAG magnetic M2 beads (M8823 Sigma) for 3 hours. Beads were washed four times with wash buffer buffer (50mM TRIS pH 7.4, 150 mM NaCl, 20 mM MgCl_2_, 0.1% wt/vol digitonin, 1x cOmplete protease inhibitor). Samples were eluted with Laemmli buffer and analyzed by immunoblot.

### Antibodies

The following antibodies were used: anti-SPTLC1 (Sigma HPA063907), anti-SPTLC2 (Abcam ab236900), anti-ORMDL3 (Millipore-SIGMA ABN417), mouse anti-FLAG (immunofluorescence, Sigma-Aldrich F1804), rabbit anti-FLAG (immunoblot, Proteintech 20543-1-AP) anti-VAPB (Proteintech 14477-1-AP), anti-UBB (Cell Signaling 3933), anti-beta-ACTIN (GenScript A00702), anti-TOMM40 (Proteintech 18409-1-AP), anti-HSPA5/BiP (Abcam ab21685); and the secondary antibodies anti-mouse Alexa Fluor 488 (Invitrogen A-11029), anti-mouse-HRP (Jackson ImmunoResearch 115-035-146), anti-rabbit-HRP (Jackson ImmunoResearch 111-035-003).

### Silencing of ORMDL and SPTLC1-ALS variants

siRNAs targeting human ORMDL1, -2 and -3 (40) were used to silence ORMDL expression. The allele specific silencing of the mutants L39del and F40S41del was based on a previously reported siRNA approach (4). For HEK293 cells, 10 nM of each siRNA was co-transfected. For Fibroblasts, 10 nM of the variant targeting siRNA was transfected. The siRNAs were diluted in reduced serum media (Opti-MEM; Gibco, 31985-047). Transfection was performed using Lipofectamine RNAiMAX Transfection Reagent (Thermo Fisher, 13778030) according to manufacturers instructions. The media was replaced after 4 hours with fresh growth media (DMEM / 10% FBS) and cells were allowed to grow for 72 hours before the labelling experiments.

### Stable isotope labelling assay in cells

Sphingolipid labelling assays and SPT activity measurements were performed in vivo in cells. SPTLC1-KO cells expressing SPTLC1-ALS and -HSAN1 variants were plated at 200,000 cells per ml in 6 well plates. Cells were grown for 48 hours to 70% confluence in DMEM 10% FBS growth media. For standard labelling assay, the media was exchanged to L-serine free DMEM (Genaxxon Bioscience, Ulm, Germany) containing 10 % FBS, 1 % P/S and isotope-labelled D_3_-^15^N-L-serine (1 mM unless indicated otherwise) and D_4_-L-alanine 2 mM (Cambridge Isotope Laboratories, MA, USA). Cells were grown further in the labelling media for 16 hours. C_6_-Cer, when used in activity assays, was added together with the labelling media. For siRNA knockdown in HEK293 cells and fibroblasts, the assay was started 72 hours post transfection. For lipid analysis, HEK293 cells were harvested in ice cold PBS. Fibroblasts were harvested by trypsinization (500 µl). Finally, 50 µL cells were counted (Z2 Coulter counter, Beckman Coulter, CA, USA) and cell pellets were frozen at -20°C until extraction.

### Amino acid analysis

Amino acids were quantified from 10 µL plasma precipitated with 180 µl ice-cold methanol containing 1 nmol of stable isotope labelled amino acids (Cambridge Isotope Laboratories, MSK-A2-1.2). Samples were incubated at -20°C for 30 min followed by centrifugation at 4°C (14,000g, 10 min). The supernatant was transferred to a fresh tube and dried under a N_2_ stream, and stored at -20°C until analysis. Dried pellets were re-constituted in 100 µL of 0.1 % acetic acid. The dissolved material was transferred to an autosampler vials following centrifugation at RT (16,000g, 5 min). Amino acids were separated on a reverse-phase C18 column (EC 250/2 NUCLEOSIL 100-3 C18HD, L=250 mm, ID: 2 mm; Macherey-Nagel). Samples (5 µL) were subjected to liquid chromatography coupled with multiple reaction monitoring (MRM) mass spectrometry using a QTRAP 6500+ LC-MS/MS-MS System (Sciex). Solvent systems used were (A) 0.1 % formic acid in water and (B) acetonitrile (100 %). The amino acids were chromatographed isocratically with solvent A for 5 minutes, followed by a linear gradient to 50 % solvent B over two minutes. Then, the column was washed with 80 % solvent B prior equilibration with 100 % solvent A. The flow rate was held constant at 0.2 ml/min. Sample ionization was achieved via electrospray ionization in positive ion mode. Quantification was performed using MultiQuant (2.1) software (SCIEX).

### Sphingolipidomics

Frozen cell pellets were resuspended in 50 μl PBS and extracted with 1 ml Methanol/MTBE/Chloroform (MMC) (4/3/3) containing 100 pmoles each of D_7_SA (d18:0), D_7_SO (d18:1), dhCer (d18:0/12:0), ceramide (d18:1/12:0), glucosylceramide (d18:1/8:0), SM (d18:1/18:1(D_9_)), and D_7_-S1P. Lipids were extracted continuously a Thermomixer (Eppendorf) at 37°C (1400 rpm, 60 minutes). The single-phase supernatant was collected, dried under N_2_ and dissolved in 100 μl methanol. Untargeted lipid analysis was performed on a high-resolution Q-Exactive MS analyser (Thermo Scientific) after lipids were separated by liquid chromatography carried out according to (2) with some modifications. Lipids were separated using a C30 Accucore LC column (150 mm x 2.1 mm, 2.6 µm particle size) and a Transcend UHPLC pump (Thermo Fisher Scientific). Liquid chromatography was performed with solvents, acetonitrile:water (6:4) with 10 mM ammonium acetate and 0.1 % formic acid, B) isopropanol:acetonitrile (9:1) with 10 mM ammonium acetate and 0.1 % formic acid at a flow rate of 0.260 ml/min. The following gradient was applied; **1**. 0.0 - 0.5 min; 70 % A, 30 % B, **2**. 0.5 - 2.0 min (57% A, 43% B), **3**. 2.0 - 3.30 min, 45 % A, 55 % B, **4**. 3.30 - 12.0 minutes 25 % A, 75% B), **5**. 12.0-18.0 100% B, **6**. 18.0-25.00 100 % B, **7**. 25.0 - 25.50 70 % A, 30 % B, **8**.0 25.50-29.50 70 % A, 30%. Lipids were similarly extracted and quantified from 50 ul patient or control plasma. MS2 fragmentation was based on data dependent acquisition (DDA). Lipid identification criteria were: (i) resolution with an accuracy of 5 ppm from the predicted mass at a resolving power of 70’000 at 200 m/z, (ii) isotopic distribution, (iii) fragmentation pattern and (iv) expected retention time relative to internal and external standards. Similar criteria were applied to isotope labelled de novo produced sphingolipids in cells carrying additional +3 Da.

Transitions used for labelled and non-labelled lipids;

1. Ceramide: [M+H]^+^ → [M+H - H_2_O]^+^, [M+H - 2H_2_O]^+^, [M+H - H_2_O - FA]^+^, and [M+H - 2H_2_O - FA]^+^
2. 1-Deoxyceramide: [M+H]^+^ → [M+H - H_2_O]^+^ and [M+H - H_2_O -FA]^+^
3. Hexosylceramide: [M+H]^+^ → [M+H - H_2_O]^+^, [M+H - 2H_2_O]^+^, [M+H - H_2_O - FA - Hex]^+^, [M+H - 2H_2_O - FA - Hex]^+^
4. Sphingomyelin: [M+H]^+^ → [M+H - H_2_O]^+^, [M+H - 2H_2_O]^+^, [M+H - H_2_O - FA - PO_4_Choline]^+^, [M+H - 2H_2_O - FA - PO_4_Choline]^+^
5. Sphingoid base (d181, d18:0): [M+H]^+^ → [M+H - H_2_O]^+^ and [M+H - H_2_O - H_2_O]^+^
6. 1-deoxysphingoid base (m18:1, m18:0): [M+H]^+^ → [M+H - H_2_O]^+^

### Statistical analysis

If not indicated differently, data were compared by one-way ANOVA with Bonferroni adjustment for multiple comparisons. Plasma SL (Supplemental Fig 3) and 1-deoxySL (Fig 6B) were compared using 2-way ANOVA with Dunnett’s adjustment for multiple comparisons. All results are expressed as mean ± standard deviation (SD). Adjusted p-value <0.05 was considered statistically significant. Statistical analyses were performed with GraphPad Prism 8.0 (GraphPad Software, Inc., San Diego, CA) and MetaboAnalyst v5.0 (www.metaboanalyst.ca) (41).

## Acknowledgments

We are grateful to the patients and their families.

## Data Availability Statement

This study did not generate any new code.

The raw data that support the findings of this study are available from the corresponding authors upon request.

