## Supplementory Fig 1, Supplementory Fig 2, Supplementory Fig 3, Supplementory S4, Supplementsry Fig 5, Supplementary Fig 6, Supplementary Fig 7 for "SPTLC1 variants associated with childhood onset amyotrophic lateral sclerosis produce distinct sphingolipid signatures through impaired interaction with ORMDL proteins"

**Legends to supplementary figures:**

**Figure S1. The de novo sphingolipid synthesis pathway.**

(**A**) Sphingolipid synthesis starts with the formation of aliphatic amino alcohols or long chain bases (LCB), the identifying structural units of all SLs. This reaction is carried out by the serine-palmitoyltransferase (SPT) using palmitoyl-CoA and L-serine. Therefore, majority of SL in mammals are composed of canonical C18 LCBs, sphingosine (d18:1) and sphinganine (d18:0). Dihydroceramide synthases CerS1-6 conjugate LCB with activated fatty acids (acyl-CoA's) of variable chain lengths ($C_{16}$ to $C_{26}$) to form dihydroceramide (dhCer, d18:0) species. Desaturases, DEGS1 and FADS3 introduce double bonds at C4-C5 and C14-C15 positions in LCB of dhCer to form mono- (d18:1) and di-unsaturated (d18:2) ceramides, respectively. Ceramides are converted to complex sphingolipids, such as sphingomyelin and glycosphingolipids with addition of specific head group structures. In a catabolic pathway, ceramides are degraded back to release LCB sphingosine (SO) and fatty acid by ceramidases. Phosphorylation of the primary hydroxyl group by sphingosine kinases SK1/2 lead to formation of a sphingosine-1-phosphate (S1P) which is degraded terminally by S1P lyase SGPL1.

(**B**) Schematic representation of SPTLC1-ALS and -HSAN1 variants relative to the ER luminal and cytosolic side.

**Figure S2. ORMDLs associate with SPT-complex.**

**(A)** SPTLC1, SPTLC2 and ORMDL protein levels in blue native PAGE samples from Fig 2B. Samples from Fig 2B were analyzed on a denaturing SDS-PAGE and immunoblotted with anti-SPTLC1, anti-SPTLC2 and anti-ORMDL antibodies. TOMM40 was used as a loading control. * = carryover signal from SPTLC2 antibody.

**(B-C)** ORMDL-protein levels in SPTLC1-KO and SPTLC2-KO cells. Whole cell lysates from Flp-In T-REx 293 control, SPTLC1-KO and SPTLC2-KO cells were analyzed by SDS-PAGE and immunoblotting (**B**), and ORMDL levels were quantified (**C**). ORMDL signals were normalized to ACTIN signal. Mean ± SD, n=4 independent replicates, unpaired two-sided Welch's t-test, *** $p<0.001$

**(D)** SPTLC1, SPTLC2 and ORMDL protein in native PAGE samples from Fig 2C. Samples from Fig 2C were analyzed on denaturing SDS-PAGE and immunoblotted with anti-SPTLC1, anti-SPTLC2 and anti-ORMDL antibodies. TOMM40 was used as a loading control.

(**E-F**) Analysis of SPT complex in patient fibroblasts by blue native PAGE. Membrane fractions from immortalized control fibroblasts (3 different control cells) and patient fibroblasts derived from S331Y and L39del patients were analyzed by blue native PAGE and immunoblotted with anti-SPTLC1, anti-SPTLC2 and anti-ORMDL antibodies **(E)**. Protein levels in these samples were analyzed on a denaturing SDS-PAGE **(F)** where TOMM40 and HSPA5 were used as loading controls. * = carryover signal from SPTLC2 antibody.

**Figure S3. Plasma sphingolipid levels in ALS and HSAN1 patients**

**(A-D)** Circulating sphingolipid levels in ALS patients, carrying SPTLC1p.Y23F, L39del and F40S41del mutations. Total ceramides (**A**), total sphingomyelins (**B**), total monohexosyleramides (**C**) and 1-deoxysphingolipids (**D**). n=4 (independent replicates), Data are represented as mean ± SD, two-way ANOVA with Dunnett's adjustment for multiple comparisons. * $p< 0.05$, *** $p<0.001$, **** $p<0.0001$.

**Figure S4. Sphingolipid species levels in ALS and HSAN1 variant expressing cells.**

(**A-B**) Levels of indicated de novo formed dihydroceramides (dhCer, **A**) and ceramides (Cer, **B**) with conjugated intermediate chain length fatty acids. HEK293 control and variant expressing SPTLC1-KO cells were grown in the presence of $D_3$, $^{15}$N-L-serine and $D_4$-L-alanine to label SLs. Absolute levels of each SL species was quantified relative to internal lipid standard by high resolution mass spectrometry. Data are represented as mean ± SD, n=3 independent replicates, one-way ANOVA followed by Bonferroni adjustment for multiple comparison, * $p< 0.05$, ** $p<0.01$, *** $p<0.001$, **** $p<0.0001$.

**Figure S5. Sphingolipid species profiles in patient derived primary fibroblasts carrying SPTLC1-ALS disease variants.**

(**A** and **B**) Dihydroceramide (**A**) and ceramide (**B**) profiles based on conjugated fatty acids (FA) in control and patient derived primary fibroblasts carrying SPTLC1p.L39del and F40S41del mutations. Cells were grown in presence of $D_3$, $^{15}$N-L-serine and $D_4$-L-alanine for 16 hours.

**(C)** Ceramide species profiles based on conjugated fatty acids (FA) in patient derived fribroblasts after transfection with scramble and mutant specific siRNA and in comparison to scramble transfected control cells. Isotope labelling was performed 72 hours post SiRNA (10 nM) transfection.

Data are represented as mean ± SD, n=3 and one-way or two-way ANOVA followed, respectively, by Bonferroni or Dunnet's adjustment for multiple comparison, * $p< 0.05$, ** $p<0.01$, *** $p<0.001$, **** $p<0.0001$.

**Figure S6. Plasma ceramide species profiles from patients carrying SPTLC1-ALS disease variants.**

(**A** and **B**) Dihydroceramide (**A**) and ceramide (**B**) profiles based on conjugated fatty acids (FA) in plasma of healthy controls, ALS patients carrying SPTLC1p.Y23F, L39del and F40S41del mutations and HSAN1 C133W patients. n=4 (independent replicates). Data are represented as mean ± SD, two-way ANOVA with Dunnett's adjustment for multiple comparisons. * $p< 0.05$, *** $p<0.001$, **** $p<0.0001$.

**Figure S7. Plasma L-serine and L-alanine levels in Leu39del family.**

(**A** and **B**) Levels of L-serine (**A**) and L-alanine (**B**) in plasma of SPTLC1p.Leu39del family members. Amino acids were extracted from plasma and quantified relative to isotope labelled internal standards. Data are represented as mean ± SD, n=4 (independent replicates), one-way ANOVA followed by Bonferroni correction for multiple comparison, * $p< 0.05$, *** $p<0.001$, **** $p<0.0001$.

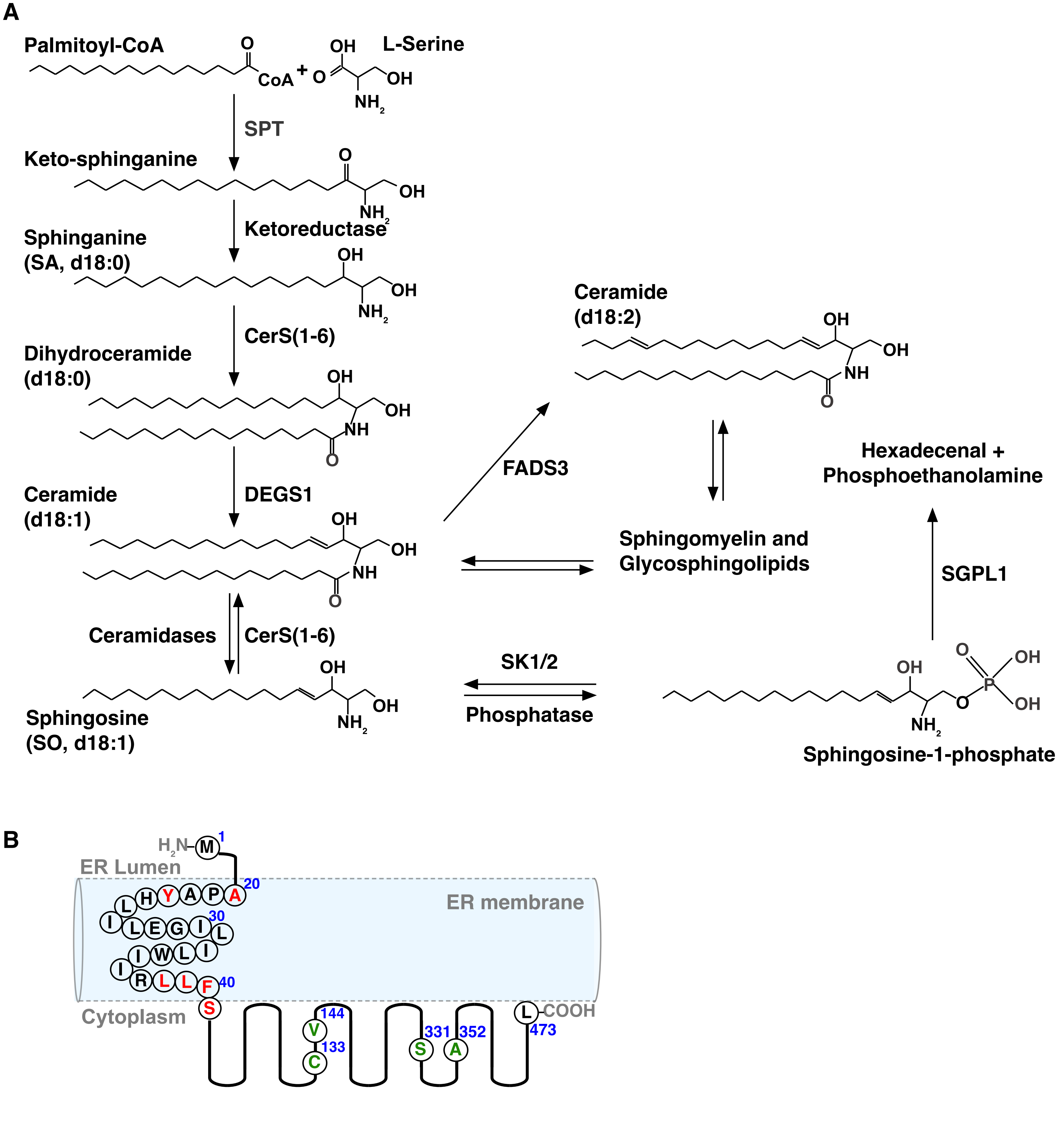
A
Palmitoyl-CoA
L-Serine
CoA
OH
NH2
SPT
Keto-sphinganine
Ketoreductase
Sphinganine
(SA, d18:0)
CerS(1-6)
Dihydroceramide
(d18:0)
NH
DEGS1
Ceramide
(d18:1)
Ceramidases
CerS(1-6)
Sphingosine
(SO, d18:1)
FADS3
Ceramide
(d18:2)
Hexadecenal +
Phosphoethanolamine
Sphingomyelin and
Glycosphingolipids
SGPL1
SK1/2
Phosphatase
Sphingosine-1-phosphate
B
H2N
ER Lumen
ER membrane
Cytoplasm
COOH
1
20
30
40
144
133
331
352
473

### Figure S1

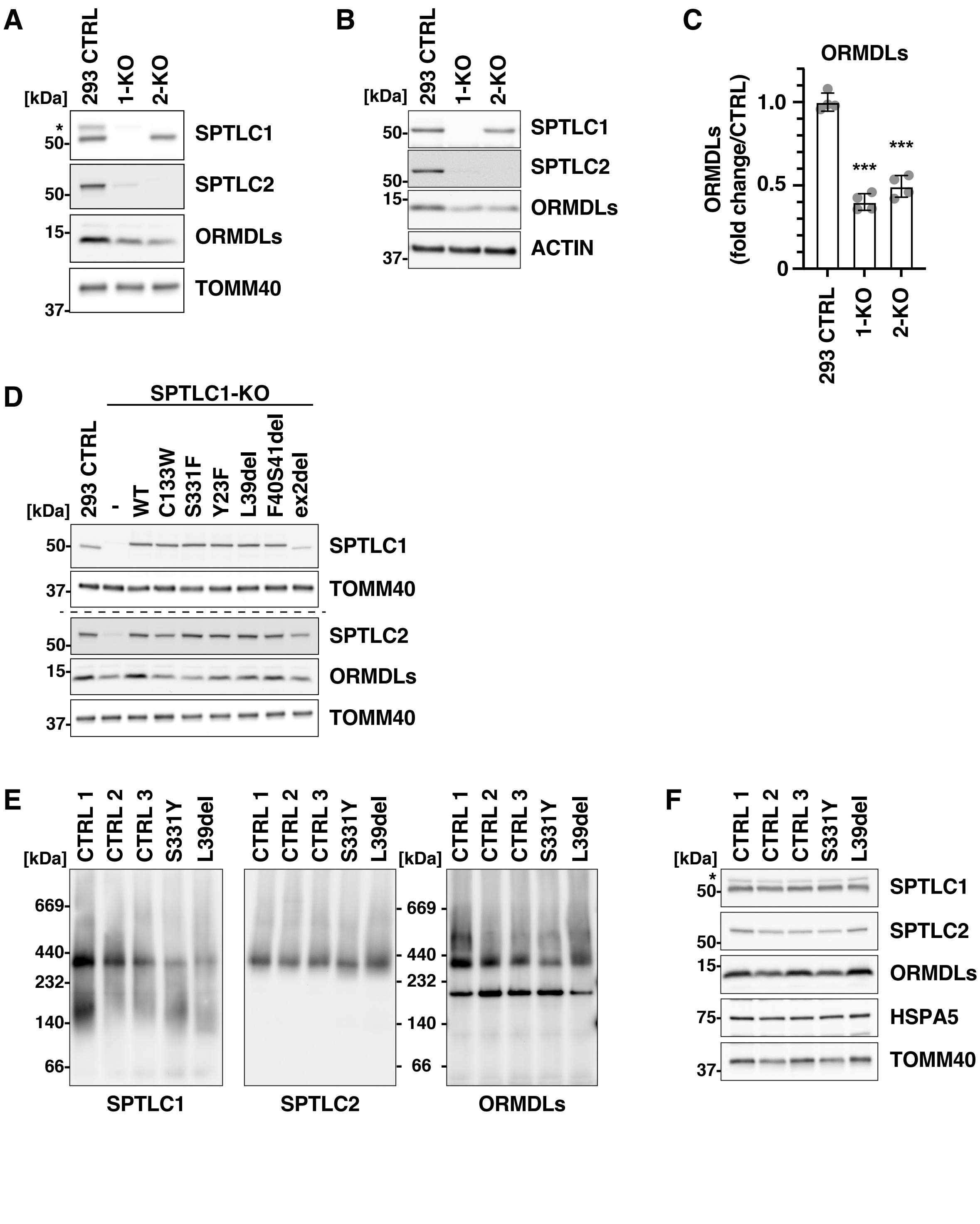
A
293 CTRL
1-KO
2-KO
[kDa]
SPTLC1
SPTLC2
ORMDLs
TOMM40
B
ACTIN
C
ORMDLs
ORMDLs (fold change/CTRL)
1.0
0.5
0
***
D
SPTLC1-KO
-
WT
C133W
S331F
Y23F
L39del
F40S41del
ex2del
E
CTRL 1
CTRL 2
CTRL 3
S331Y
669
440
232
140
66
F
HSPA5

### Figure S2

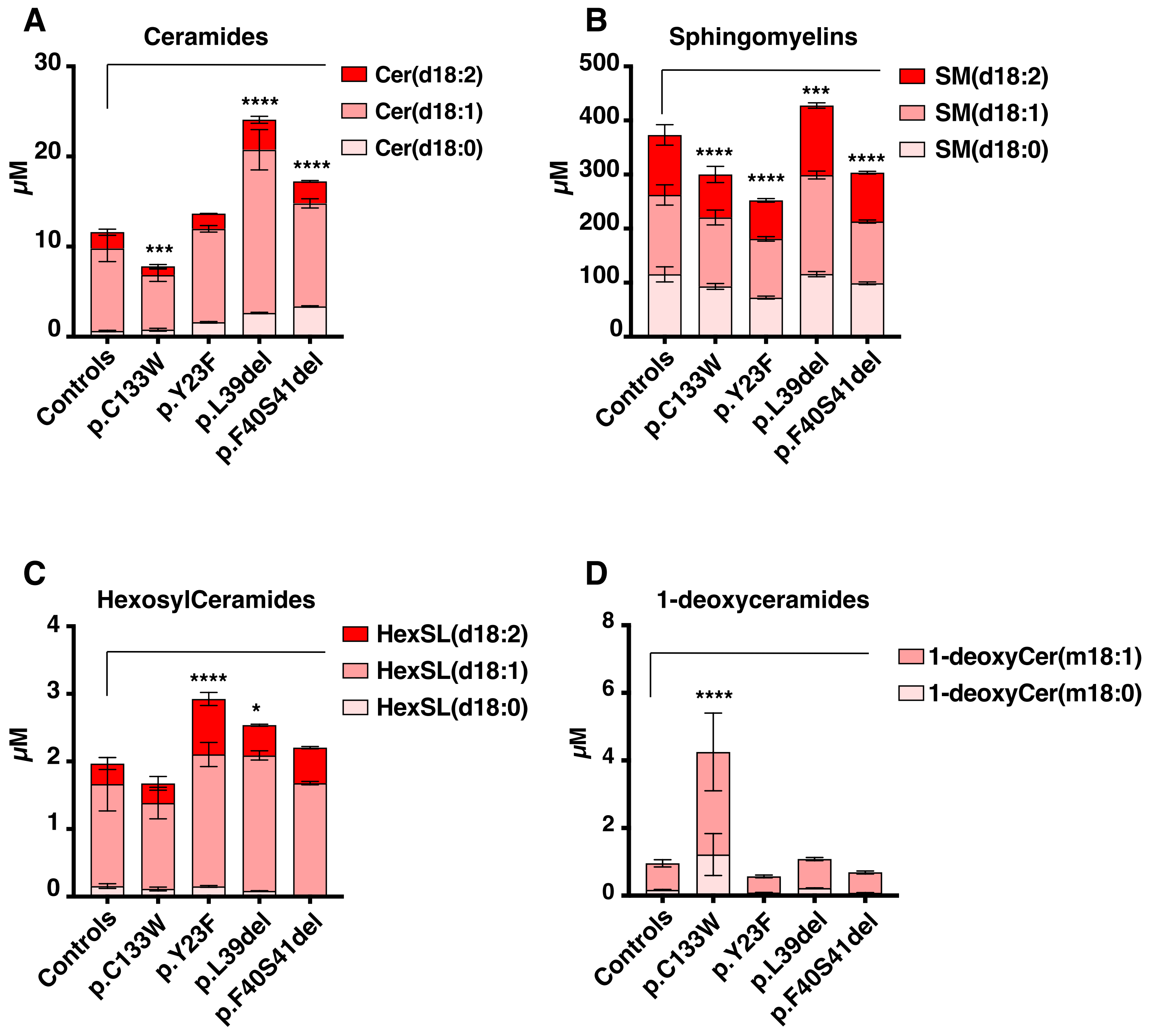
A
Ceramides
Cer(d18:2)
Cer(d18:1)
Cer(d18:0)
μM
30
20
10
0
***
****
****
Controls
p.C133W
p.Y23F
p.L39del
p.F40S41del
B
Sphingomyelins
SM(d18:2)
SM(d18:1)
SM(d18:0)
μM
500
400
300
200
100
0
****
****
***
****
Controls
p.C133W
p.Y23F
p.L39del
p.F40S41del
C
HexosylCeramides
HexSL(d18:2)
HexSL(d18:1)
HexSL(d18:0)
μM
4
3
2
1
0
****
*
Controls
p.C133W
p.Y23F
p.L39del
p.F40S41del
D
1-deoxyceramides
1-deoxyCer(m18:1)
1-deoxyCer(m18:0)
μM
8
6
4
2
0
****
Controls
p.C133W
p.Y23F
p.L39del
p.F40S41del

### FIGURE S3

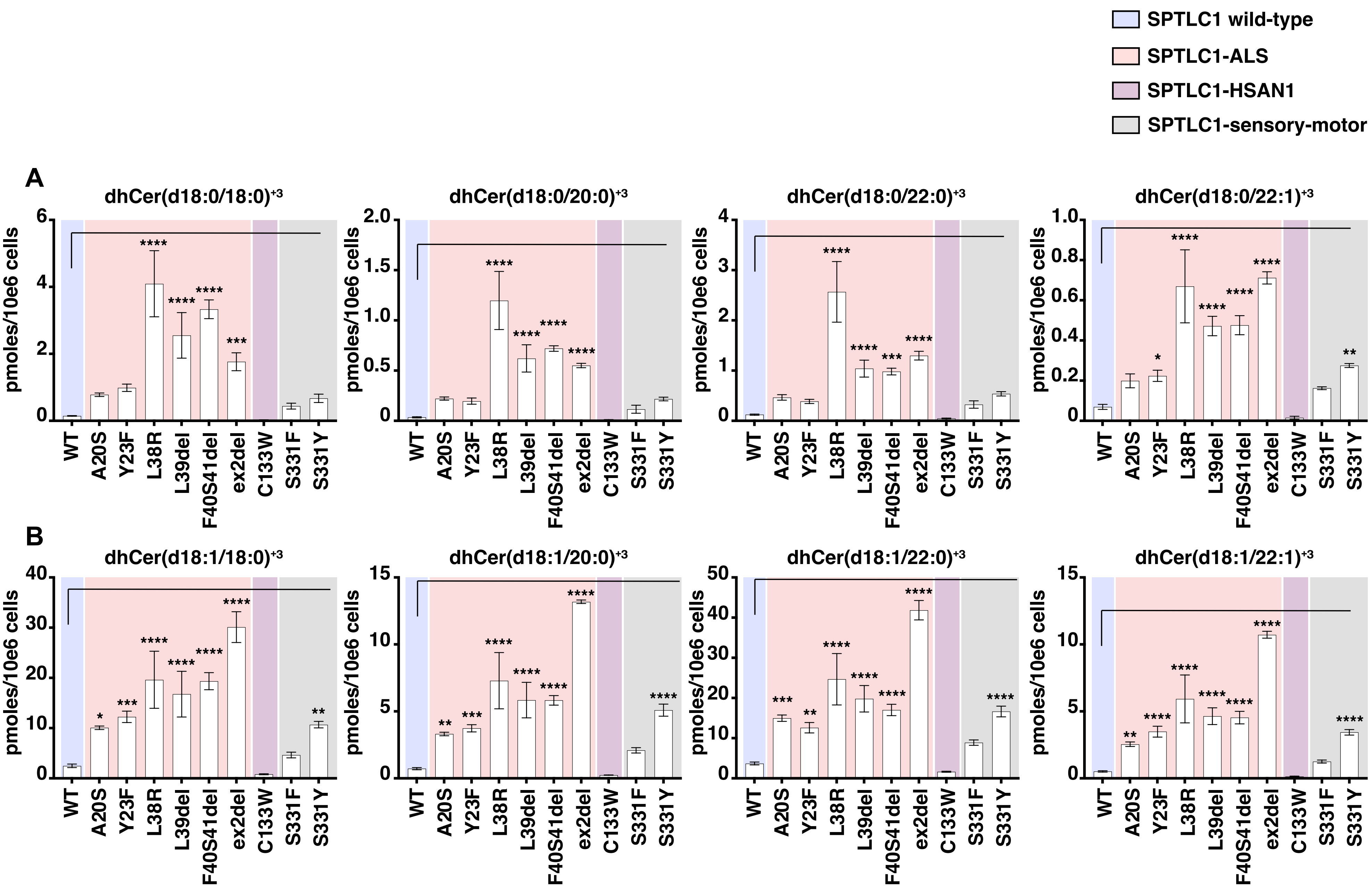
SPTLC1 wild-type
SPTLC1-ALS
SPTLC1-HSAN1
SPTLC1-sensory-motor
A
dhCer(d18:0/18:0)+3
dhCer(d18:0/20:0)+3
dhCer(d18:0/22:0)+3
dhCer(d18:0/22:1)+3
B
dhCer(d18:1/18:0)+3
dhCer(d18:1/20:0)+3
dhCer(d18:1/22:0)+3
dhCer(d18:1/22:1)+3
pmoles/10e6 cells
WT
A20S
Y23F
L38R
L39del
F40S41del
ex2del
C133W
S331F
S331Y

### Figure S4

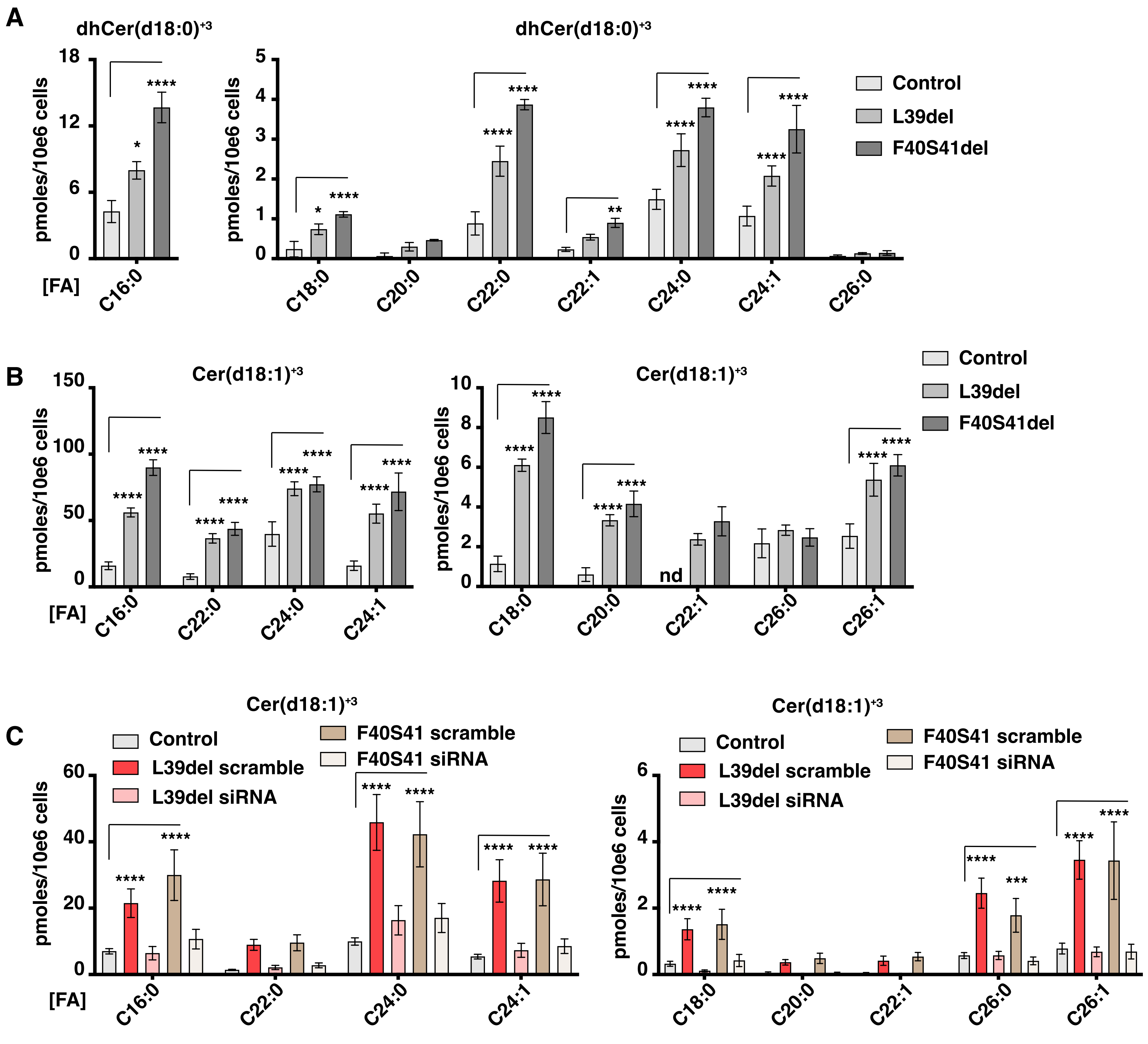
A
dhCer(d18:0)+3
pmoles/10e6 cells
[FA]
C16:0
C18:0
C20:0
C22:0
C22:1
C24:0
C24:1
C26:0
Control
L39del
F40S41del
B
Cer(d18:1)+3
nd
C
L39del scramble
L39del siRNA
F40S41 scramble
F40S41 siRNA
C26:1

### Figure S5

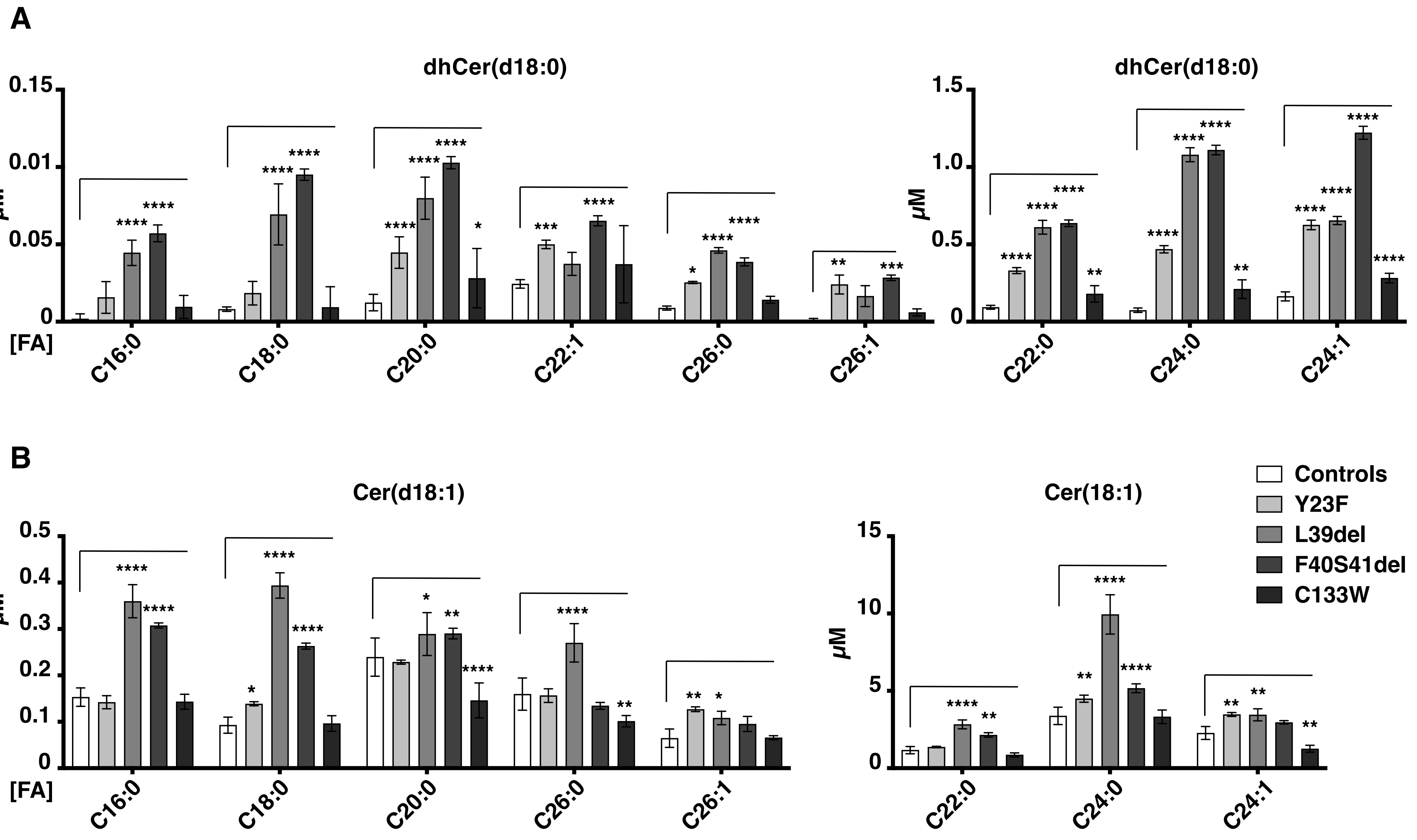
A
dhCer(d18:0)
dhCer(d18:0)
µM
[FA]
C16:0
C18:0
C20:0
C22:1
C26:0
C26:1
C22:0
C24:0
C24:1
B
Cer(d18:1)
Cer(18:1)
Controls
Y23F
L39del
F40S41del
C133W
µM
[FA]
C16:0
C18:0
C20:0
C26:0
C26:1
C22:0
C24:0
C24:1

### Figure S6

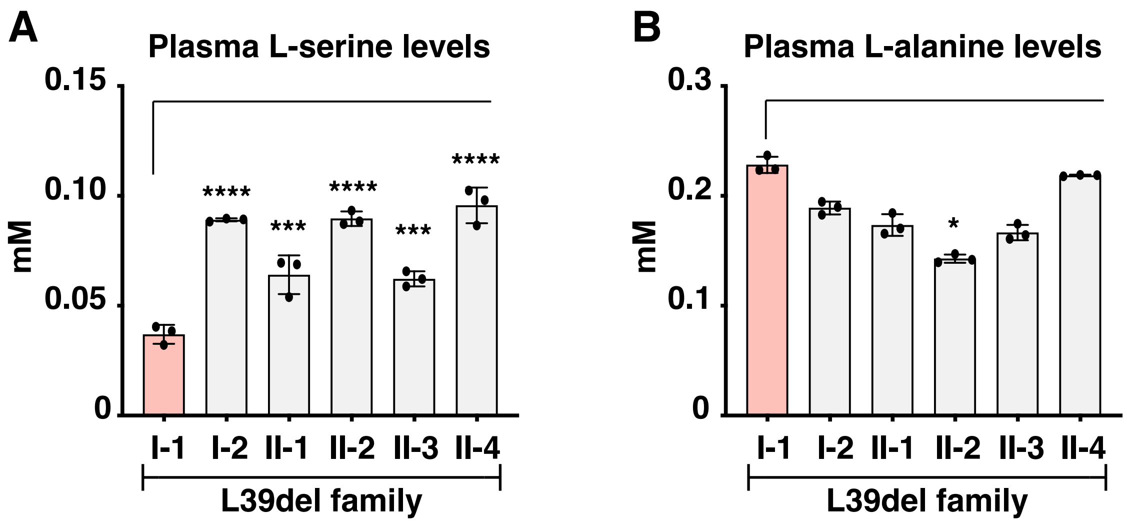
A
Plasma L-serine levels
0.15
0.10
0.05
0
mM
****
***
****
***
****
I-1
I-2
II-1
II-2
II-3
II-4
L39del family
B
Plasma L-alanine levels
0.3
0.2
0.1
0
mM
*
I-1
I-2
II-1
II-2
II-3
II-4
L39del family

### Figure S7
